# NADH dehydrogenase reverses dietary and clock metabolic syndrome

**DOI:** 10.1101/2025.10.22.684030

**Authors:** Chelsea Hepler, Nathan J. Waldeck, Benjamin J. Weidemann, Biliana Marcheva, You-Jia Chen, Jacqueline Hecker, Ziming Zhu, Rino Nozawa, Joseph V. Mastroni, Anneke K. Thorne, Colleen R. Reczek, Jonathan Cedernaes, Kathryn M. Ramsey, Clara B. Peek, Grant D. Barish, Navdeep S. Chandel, Joseph Bass

## Abstract

Circadian clocks are internal timing systems that enable organisms to anticipate and adapt to daily environmental changes. These rhythms arise from a transcription-translation feedback loop in which CLOCK/BMAL1 regulate the expression of thousands of genes, including their repressors PER/CRY^1^. Disruption of circadian rhythms contributes to obesity, metabolic disease, and cancer^2-4^, yet how the clock maintains metabolic homeostasis remains limited. Here we report that the clock regulates oxidative metabolism through diurnal respiration of mitochondrial respiratory chain complex I. Genetic loss of the clock and high fat diet feeding in male mice led to reduced complex I respiration within adipocytes, leading to suppression of PPAR and insulin signaling pathways. In contrast, preserving complex I function maintained adipogenic and metabolic gene networks and protected against diet- and circadian-induced metabolic dysfunction independently of weight gain. These findings reveal that circadian disruption impairs metabolic health through mitochondrial complex I dysfunction, establishing clock control of complex I as a key regulator of transcriptional and metabolic homeostasis.

## Main Text

High-fat feeding leads to circadian disruption^5^, suggesting that an understanding of how clock transcription cycles control physiology will elucidate general mechanisms of metabolic disease. At the molecular level, the core clock autoregulatory transcription-translation feedback loop is driven by the activators CLOCK/BMAL1 which induce the expression of the repressors PER/CRY1^1^. In the negative limb of the clock, PER/CRY then bind to CLOCK/BMAL1 to inhibit their own expression. A second loop mediated by REV-ERB/ROR stabilizes expression of *Bmal1.* While there is extensive knowledge on the interplay of core clock proteins in the central pacemaker cells driving rhythms in the sleep-wake cycle, our understanding of how peripheral clocks regulate metabolism and health remains limited.

The circadian clock plays a pivotal role in physiological rhythms through direct transcriptional control of key rate-limiting enzymes in intermediary metabolism best characterized in liver^6,7^. The clock also indirectly influences metabolism through control of epigenetic pathways important in the flexibility of fuel switching between oxidative and reductive metabolism across the fasting/feeding cycle. For example, BMAL1 directly controls transcription of the rate-limiting enzyme involved in NAD+ biosynthesis^8^. This in turn drives rhythms in fatty acid oxidation through time-dependent activation of NAD+-dependent enzymes, including the sirtuins and the poly (ADP-ribose) polymerases^9,10^. REVERBs also represent a key node linking core circadian and metabolic gene transcription^11^. Peripheral clocks also exhibit distinct tissue-specific properties; for instance, the liver clock is entrained by feeding time, whereas the adipose clock aligns with the light/dark cycle^5,12-14^.

Obesity is associated with pronounced disruption in circadian clock gene expression within adipose tissue^5,15^. Mice with genetic ablation of the adipocyte clock display enhanced weight gain during high fat diet (HFD) feeding and increased visceral adiposity^16^. These mice also exhibit arrhythmic food intake, with increased feeding during the light (rest) period^16^. These findings reveal that circadian disruption in adipose tissue uniquely contributes to systemic impairment in metabolic health during overnutrition. Our prior work revealed a role for the adipocyte circadian clock in regulating energy balance according to the time when an animal eats through control of creatine-mediated thermogenesis^13^. Mice lacking the circadian clock in adipocytes have reduced creatine abundance and futile creatine cycling in adipose tissue. However, genetic ablation of the circadian clock in adipocytes exerts a significant impact on metabolites and enzymes extending well beyond the scope of creatine thermogenesis^17,18^. Therefore, we sought to examine how the circadian clock governs bioenergetics in adipocytes.

## Results

### The circadian clock regulates respiration at mitochondrial complex I

To determine how the circadian clock impacts mitochondrial metabolism in adipose tissue, we performed a comprehensive assessment of mitochondrial electron transport chain (ETC) function at different times of the subjective day and night. We isolated mitochondria from visceral white adipose tissue (WAT) in mice entrained to a normal or inverted light cycle to compare mitochondria from mice at different zeitgeber times (ZT). Mitochondrial oxygen consumption rate (OCR) in the presence of the complex I (CI)-linked substrate pyruvate was elevated during the dark/active period (ZT14) as compared to the light/inactive period (ZT2) (**Fig. 1a**). Mice lacking the clock activator *Bmal1* in adipocytes (*Bmal1*-KO) displayed reduced mitochondrial OCR in response to pyruvate during the dark period (ZT14) compared to mice with an intact clock. However, mitochondrial OCR in the presence of the complex II (CII)-linked substrate succinate was not different in circadian mutant adipose tissue mitochondria (**Fig. 1a**). Therefore, adipocyte mitochondrial respiration in response to CI (pyruvate) but not CII (succinate) substrate is diurnal and dependent on the clock.

**Figure 1.**
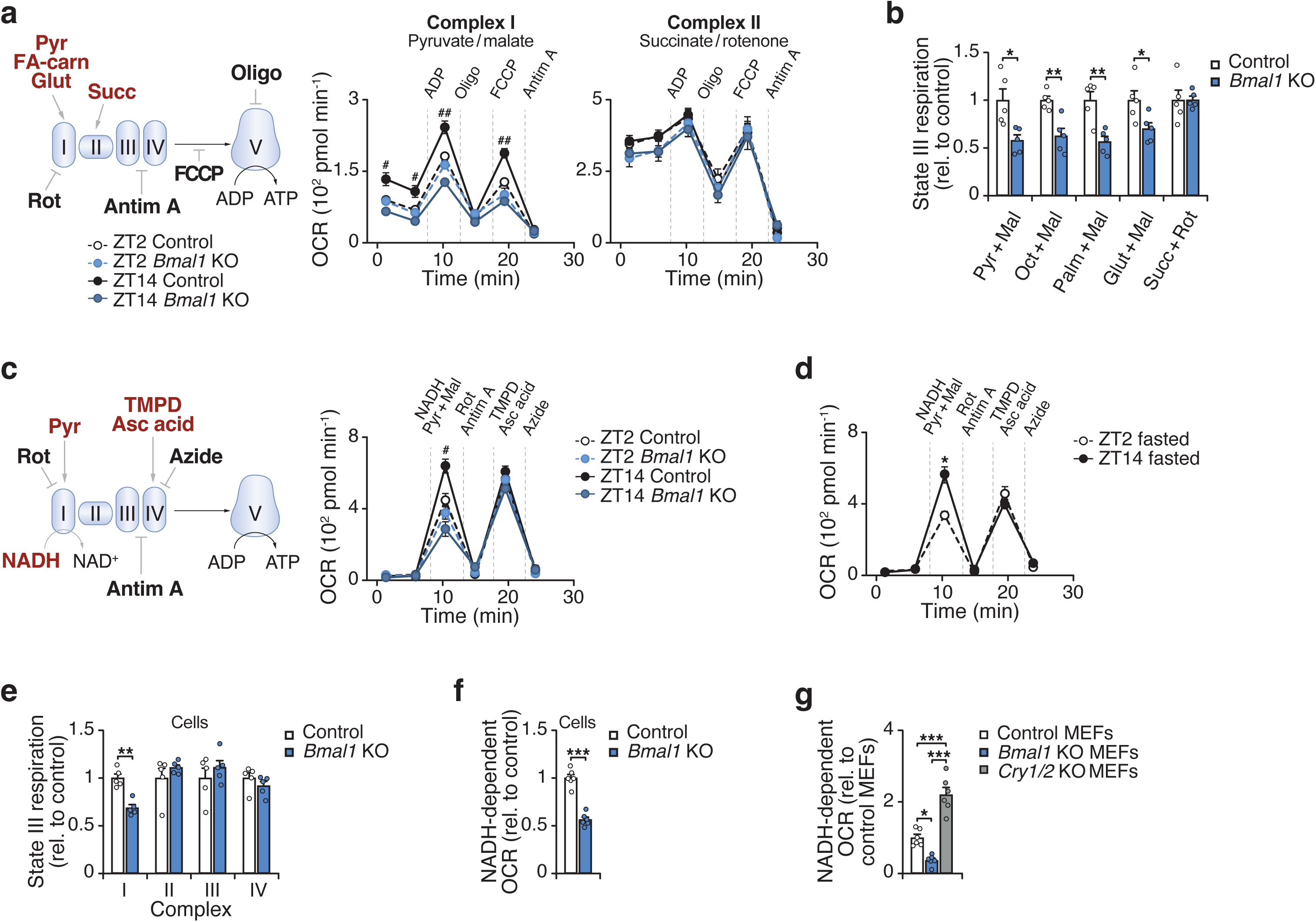
The circadian clock regulates respiration at mitochondrial respiratory chain complex I. **a**, OCR in gWAT mitochondria isolated from Control (*Bmal1^fx/fx^*) and *Bmal1*-KO (*Adiponectin-Cre; Bmal1^fx/fx^*) mice at ZT2 and ZT14 in the presence of indicated substrates and inhibitors (n = 6-7). Pyr, pyruvate; FA-carn, fatty acid carnitine; Succ, succinate; Rot, rotenone; Antim A, antimycin A; Oligo, oligomycin; FCCP, Carbonyl cyanide-p-trifluoromethoxyphenylhydrazone. **b,** State III respiration in gWAT mitochondria isolated from Control and *Bmal1*-KO mice at ZT14 in response to CI- and II-linked substrates (n = 5). **c,** OCR in gWAT mitochondria of Control and *Bmal1*-KO mice at ZT2 and ZT14 in the presence of indicated substrates and inhibitors (n = 5). TMPD, Tetramethyl-p-phenylenediamine; Asc acid; ascorbic acid. **d,** OCR in gWAT mitochondria of wildtype mice at ZT2 and ZT14 following 12 hours of fasting in response to indicated substances as in C (n = 5). **e,** State III respiration in adipocytes differentiated from Control and *Bmal*1-KO mice in the presence of CI-IV substrates (n = 5). **f,** NADH-dependent OCR in adipocytes differentiated from Control and *Bmal1*-KO mice (n=6). **g,** NADH-dependent OCR in Control, *Bmal1^-/-^* and *Cry1/2^-/-^* MEFs (n=6). Data are represented as mean ± SEM. Statistical significance was calculated by two-way ANOVA followed by Dunnett’s multiple comparisons test with ZT14 Control set as the reference group in **a** and **c**, multiple unpaired t-tests in **b** and **e**, two-way ANOVA followed by multiple comparisons in **d**, unpaired t-test in **f**, and one-way ANOVA followed by multiple comparisons in **g**. The # symbol denotes significance between ZT14 Control and all other groups. *p < 0.05, **p < 0.01, ***p < 0.001.

We next assessed respiration in response to distinct substrates that feed into CI via NADH generation from glycolysis, beta oxidation, and glutaminolysis. Mitochondria from adipocyte-*Bmal1*-knockout mice had reduced ADP-stimulated (state 3) respiration in the presence of NADH-generating substrates, but not succinate (**Fig. 1b**). To determine whether the respiratory deficit in circadian mutant adipocytes lies at or upstream of CI, we measured respiration in response to NADH. We found that NADH-fueled respiration was elevated in visceral WAT mitochondria during the dark/active period (ZT14) and reduced in adipocyte-*Bmal1*-KO mitochondria (**Fig. 1c**). We also examined a subcutaneous WAT depot (inguinal WAT) and found that CI respiration was also rhythmic and clock-dependent in this depot, consistent with the phenotype observed in gWAT (**Extended Data Fig. 1a**).

To assess whether diurnal activation of mitochondria reflects an intrinsic rhythmic property or feeding state, we analyzed WAT mitochondria at ZT2 and ZT14 following a 12-hour fast. Even under these conditions, NADH-fueled CI respiration remained higher at ZT14 than ZT2 (**Fig. 1d**). This finding indicates that diurnal CI activity persists in fasted mice. However, a single overnight fast does not fully eliminate feeding effects. Therefore, to circumvent feeding rhythms altogether, we next performed Seahorse assays in cultured adipocytes. In permeabilized adipocytes, OCR in response to CI substrate (pyruvate), but not CII (succinate), CIII (duroquinol), or CIV (TMPD/ascorbate), was reduced in *Bmal1*-deficient cells (**Fig. 1e**). NADH-fueled CI respiration was also rhythmic in synchronized adipocytes (**Extended data Fig. 1b-d**) and decreased in adipocytes lacking *Bmal1* in vitro (**Fig. 1f**). Consistent with these results, mouse embryonic fibroblasts (MEFs) lacking the clock activator *Bmal1* displayed reduced CI respiration, whereas MEFs lacking the repressors *Cry1* and *Cry2* exhibited increased CI respiration (**Fig. 1g**). These data reveal opposing effects of the forward (BMAL1-mediated) and reverse (CRY-mediated) limbs of the clock on CI respiration, independent of nutrient availability. Collectively, these data reveal that mitochondrial respiration at CI is diurnal and under intrinsic circadian control.

Mammalian mitochondrial CI is the largest respiratory enzyme, composed of 45 subunits^19^. The assembly of CI necessitates the precise coordination of various assembly factors as well as all nuclear and mitochondrial DNA-encoded subunits to form a fully operational respiratory complex. This process is regulated both transcriptionally as well as through several post-translational modifications driven by nutrient and redox status that modify CI assembly, stability, and function^20^. Ablation of the clock did not affect mRNA abundance of nuclear and mitochondrial DNA-encoded CI subunits (**Extended Data Fig. 2a**). However, the protein abundance of the CI subunit NDUFA9 was reduced in mitochondria from adipocyte-*Bmal1*-knockout mice, while NDUFA10 and NDUFS2 were trending lower but not significant (p=0.07 and p=0.1, respectively) (**Extended Data Fig. 2b**). Consistent with the loss of some protein of CI subunits, the total abundance of CI was reduced in mitochondria from adipocyte-*Bmal1*-knockout mice (**Extended Data Fig. 2c**). In addition, WAT mitochondria from mice isolated at ZT2 showed a decreased abundance of CI as compared to ZT14 (**Extended Data Fig. 2d**). Neither genetic deletion of the clock or time-of-day affected the total amount of CII-V. Together these results indicate that the abundance of mitochondrial CI is regulated by the circadian clock independently of direct transcriptional control.

### Preserving respiration at complex I in adipocytes improves metabolic health during obesity

Genetic studies in mice have shown that reduced mitochondrial function in adipocytes plays a causative role in systemic insulin resistance^21,22^. Obesity is associated with impaired adipose tissue mitochondrial function as well as a reduction in circadian clock genes in adipose tissue^5,23,24^. Supporting this finding, we observed reduced and loss of diurnal CI respiration in visceral WAT mitochondria after HFD feeding (**Extended Data Fig. 3a,b**). We hypothesized that circadian disruption during obesity drives impaired mitochondrial function and subsequent metabolic disease in part through dysregulation at CI.

To investigate the role of adipocyte CI function, we generated mice with conditional expression of yeast NADH dehydrogenase (NDI1)^25,26^ in adipocytes (*Adiponectin-Cre;NDI1^LSL^*). NDI1 is a rotenone-insensitive, non-proton translocating NADH dehydrogenase that regenerates NAD+ and transfers electrons to ubiquinone (**Fig. 2a,b**)^27^. Therefore, NDI1 can restore oxidative phosphorylation through functionally replacing CI^20,28^. In lean adult mice, the expression of NDI1 in adipocytes had no impact on body weight, glucose homeostasis, the expression of mature adipocyte and thermogenic markers, or histological features of adipose depots (**Extended Data Fig. 4a-e**).

**Figure 2.**
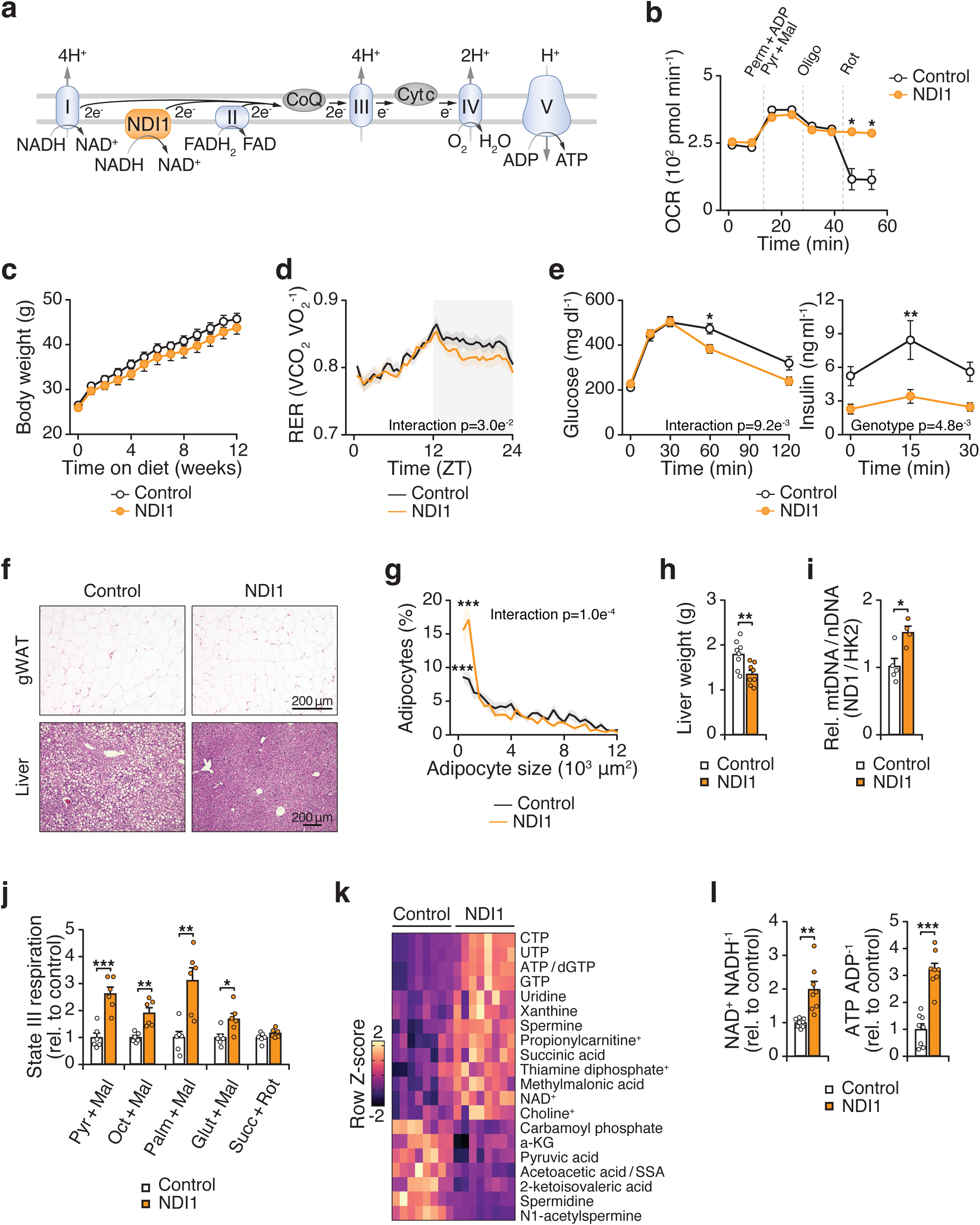
Expression of the yeast NDI1, an alternative NADH dehydrogenase, in adipocytes improves metabolic health during high fat diet feeding. **a**, Schematic of the mitochondrial electron transport chain with ectopic yeast NDI1. NDI1 transfers electrons to ubiquinone and regenerates NAD+. **b,** OCR of adipocytes differentiated from Control (*NDI1^LSL^*) and NDI1 (*Adiponectin-Cre; NDI1^LSL^*) mice in the presence of indicated substrates and inhibitors (n = 5). **c,** Body weight during high fat diet (HFD) feeding for 12 weeks in Control and NDI1 mice (n = 8). **d,** RER over 24 hours after 10 weeks of HFD feeding (n = 8). **e,** Glucose tolerance test (GTT) and insulin during the GTT at 12 weeks of HFD feeding (n = 8). **f,** Representative images of hematoxylin and eosin staining of gWAT and liver after 12 weeks of HFD feeding (20x and 10x magnification, respectively, scale bars 200 μm). **g,** Distribution in visceral WAT adipocyte size after 12 weeks of HFD feeding (n = 5). **h,** Liver weights after 12 weeks of HFD feeding (n = 8). **i,** Mitochondria/nuclear DNA ratio in gWAT after 12 weeks of HFD feeding (n = 4-5). **j,** State III respiration in gWAT mitochondria in response to CI- and II-linked substrates after 12 weeks of HFD feeding (n = 6). **k,** Heatmap of differentially abundant metabolites (p < 0.05) in visceral WAT from Control and NDI1 mice after 12 weeks of HFD feeding at ZT14 (n = 8). **l,** The NAD+/NADH and ATP/ADP ratios in WAT from Control and NDI1 mice after 12 weeks of HFD feeding at ZT14 (n = 8). Data are represented as mean ± SEM. Statistical significance was calculated by two-way ANOVA followed by multiple comparisons in **b-f, g, and j** and unpaired t-test in **h, i** and **l**. *p < 0.05, **p < 0.01, ***p < 0.001.

We next tested whether preserving adipocyte mitochondrial CI function during HFD could prevent obesity and metabolic syndrome. Mice expressing adipocyte NDI1 displayed similar weight gain, activity, caloric intake, and energy expenditure during HFD feeding compared to control mice (**Fig. 2c**, **Extended Data Fig. 5a-c**). However, NDI1 expression led to reduced RER during the dark/active period, indicating a shift toward lipid metabolism (**Fig. 2d**). Remarkably, NDI1 expression led to improved glucose tolerance and reduced insulin release during the glucose tolerance test (**Fig. 2e**).

Histological analysis revealed smaller adipocytes in visceral WAT and reduced hepatic steatosis in mice expressing NDI1 in adipocytes (**Fig. 2f-h**). NDI1 expression did not have any major effects on gene expression in BAT and inguinal WAT (**Extended Data Fig. 5d**). However, NDI1 expression increased adipogenic markers (*Pparg2* and *Adipoq*) and decreased several inflammatory markers (*Adgre1*, *Ccl2*, *Il6*, and *Tnf*) in visceral WAT (**Extended Data Fig. 5d**). This gene expression profile, along with the smaller visceral adipocyte size, is consistent with healthy visceral WAT remodeling during HFD feeding in NDI1-expressing mice. Visceral WAT from NDI1-expressing mice exhibited increased mtDNA content and enhanced CI-but not CII-driven respiration, without changes in the abundance of either complex (**Fig. 2i,j** and **Extended Data Fig. 5e,f**). Metabolomics of visceral WAT revealed that NDI1 expression led to an increase in abundance of several nucleotides (NAD+, ATP, GTP) and succinic acid, and a reduction in pyruvic acid and alpha-ketoglutarate (**Fig. 2k**). NDI1 expression also increased the NAD+/NADH and the ATP/ADP ratios in visceral WAT (**Fig. 2l**). Thus, preserving adipocyte CI function with NDI1 sustained NADH oxidation and mitochondrial respiration, enabling visceral WAT to remodel through hyperplasia rather than hypertrophy during HFD feeding. Collectively, these findings indicate that reduced adipocyte mitochondrial CI activity during obesity is a key driver of visceral WAT metabolic dysfunction and subsequent systemic impairment in metabolic health.

### Deficiency of mitochondrial complex I in adipocytes alters adipose remodeling and systemic metabolism

The improved gene expression profile in visceral WAT in NDI1-expressing mice after HFD feeding prompted us to further examine the role of CI in visceral adipocytes. Defects in CI are among the most common causes of mitochondrial disorders and have been linked to a wide range of pathologies, including neurodegenerative and cardiac diseases^29^. Genetic CI dysfunction in various tissues remodels the global transcriptome and alters cell state through retrograde signaling^25,26^. To investigate how CI deficiency affects adipocyte biology, we generated mice lacking the core catalytic subunit of CI, *Ndufs2*, specifically within adipocytes^30^. Loss of NDUFS2 leads to complete CI deficiency since NDUFS2 encodes an essential catalytic core subunit required for enzyme assembly and electron transfer^31^.

*Ndufs2* deletion led to loss of NDUFS2 protein in isolated adipose tissue mitochondria, along with reduced CI subunit NDUFB8 and CIII subunit UQCRC2 (**Extended Data Fig. 6a-c**). Consistent with this finding, we observed less respiration in response to CI and III substrates in isolated mitochondria and primary adipocytes from mice lacking adipocyte *Ndufs2* (**Extended Data Fig. 6d,e**). Mice lacking adipocyte *Ndufs2* exhibited similar body weight to control mice on a chow diet (**Fig. 3a**). However, adipocyte *Ndufs2*-KO mice had reduced WAT mass, increased liver mass, visceral adipocyte hypertrophy, and impaired glucose tolerance (**Fig. 3b-d**). In addition, circulating fatty acids were elevated in *Ndufs2*-KO mice (**Fig. 3e**). These findings indicate that loss of CI in adipocytes comprises lipid storage capacity in adipose tissue and systemic metabolic homeostasis.

**Figure 3.**
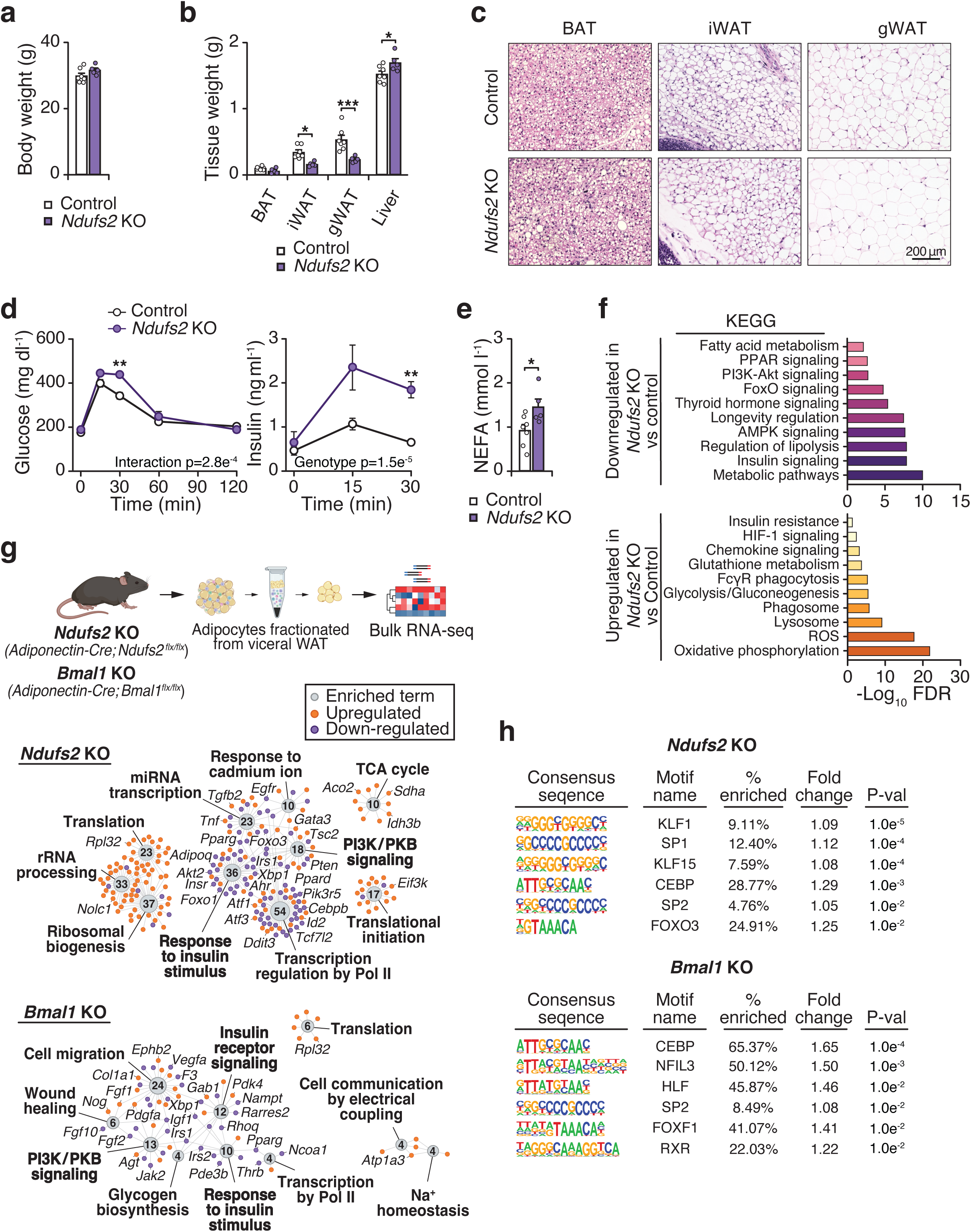
Adipocyte complex I deficiency limits adipose expansion and impairs systemic glucose homeostasis. **a**, Body weight of Control (*Ndufs2^fx/fx^*) and *Ndufs2*-KO (*Adiponectin-Cre; Ndufs2^fx/fx^*) mice at 4 months of age on a chow diet (n = 6-7). **b,** Tissue weights of Control and *Ndufs2*-KO mice at 4 months of age (n = 6-7). **c,** Representative images of hematoxylin and eosin staining of gWAT and liver 4 months of age (10x magnification, scale bars 200 μm). **d,** Oral glucose tolerance test (GTT) and insulin during the GTT at 4 months of age (n = 6-7). **e,** Serum non-esterified fatty acids (NEFA) in Control and *Ndufs2*-KO mice at 4 months of age (n = 6-7). **f,** KEGG pathway analysis of downregulated (upper panel) and upregulated (lower panel) genes in *Ndufs2*-KO vs. Control adipocytes (n = 4). **g,** RNA-sequencing was performed on fractionated adipocytes from the visceral WAT of Control (*Ndufs2^fx/fx^*), *Ndufs2-*KO (*Adiponectin-Cre;Ndufs2^fx/fx^*), and *Bmal1*-KO (*Adiponectin-Cre; Bmal1^fx/fx^*) mice. Pathway analysis of Gene Ontology biological process terms with highlighted genes among differentially expressed genes in *Ndufs2-*KO vs Control and *Bmal1*-KO vs Control fractionated adipocytes. **h,** Motif analysis of downregulated genes in *Ndufs2-*KO vs Control and *Bmal1*-KO vs Control fractionated adipocytes. Data are represented as mean ± SEM. Statistical significance was calculated by unpaired t-test in **a** and **e**, and two-way ANOVA followed by multiple comparisons in **b** and **d**. *p < 0.05, **p < 0.01, ***p < 0.001.

To define the molecular basis of the metabolic dysfunction caused by CI loss in adipocytes, we next analyzed the transcriptome in visceral WAT. We performed RNA-sequencing on the fractionated adipocytes from visceral WAT of these mice. In mice deficient in adipocyte CI, we observed reduced expression of genes involved in the response to insulin stimulus and phosphatidylinositol-3-kinase (PI3K)/protein kinase B (PKB) signal transduction (**Fig. 3f,g** and **Extended Data Table 1**). Expression of *Akt1/2*, *Irs1/2*, and *Insr*, as well as several transcription factors important in adipocyte metabolism and function such as *Pparg*, *Foxo1*, *Cebpb*, and *Tcf7l2* were reduced during *Ndufs2* deletion. In addition, we detected an upregulation in the expression of the proinflammatory cytokine *Tnf* and genes involved in the integrated stress response (*Atf3*, *Atf5*, *Ddit3*, *Dele1*), and reduced expression of the adipokine *Adipoq*. This transcriptional profile during CI dysfunction suggests a broader impairment in adipocyte metabolism and function, extending beyond isolated electron transport chain disruption.

We next compared the gene expression profile between adipocytes deficient in CI (Ndufs2-KO) and adipocytes lacking the clock (*Bmal1*-KO). Genetic deletion of the clock activator *Bmal1* led to reduced expression of genes involved in biological processes of insulin receptor signaling and PI3K/PKB signal transduction (*Irs1/2, Pdk2, Igf1*) and transcription (*Pparg, Ncoa1, Thrb*) (**Fig. 3g**, **Extended Data Table 1**). Both genetic deletion of CI and the clock resulted in a downregulation of genes involved in the insulin response and mRNA transcription, and an increase in genes involved in translation. Among the overlapping downregulated genes were *Pparg* and *Cebpa*, master regulators of fat cell metabolism, insulin sensitivity, and adipogenesis, and *Ppargc1b*, a transcriptional coactivator for PPARs^32^. We also noted an increase in the integrated stress response genes *Atf5* and *Dele1* and several CI subunits (*Ndufab1*, *Ndufb9*, *Ndufs4*, *Ndufs5*, *Ndufv2*). Motif analysis of downregulated genes in both *Ndufs2*- and *Bmal1*-KO adipocytes revealed shared enrichment for CEBP and SP family transcription factor motifs (**Fig. 3h**). These factors are key regulators of *Pparg* expression and adipogenic gene networks, consistent with the observed transcriptional changes. The shared depletion of CEBP and SP motifs suggests that disruption of CI and circadian clock function converge to maintain adipogenic transcriptional identity and insulin-responsive gene expression.

### NDI1 expression prevents metabolic dysfunction in adipocyte circadian mutant mice

Prior work has shown that mice lacking the clock in adipocytes exhibit impaired glucose homeostasis during chow and HFD feeding^13,16^. To test whether a defect at CI underlies this phenotype, we bred mice expressing NDI1 to mice deficient in *Bmal1* in adipocytes. NDI1 expression in differentiated *Bmal1*-KO adipocytes restored the NAD+/NADH ratio and CI respiration (**Fig. 4a**). On a chow diet, adipocyte-specific loss of *Bmal1* impaired glucose tolerance and caused compensatory hyperinsulinemia despite normal body weight across groups (**Extended Data Fig. 7a,b**). Expression of NDI1 in adipocyte-Bmal1-KO mice normalized glucose tolerance and insulin secretion under these conditions. These results indicate that NDI1 rescues metabolic defects caused by clock disruption independently of dietary stress. Since CI respiration is further compromised during HFD feeding, we next asked whether NDI1 could preserve metabolic function under this added metabolic stress.

**Figure 4.**
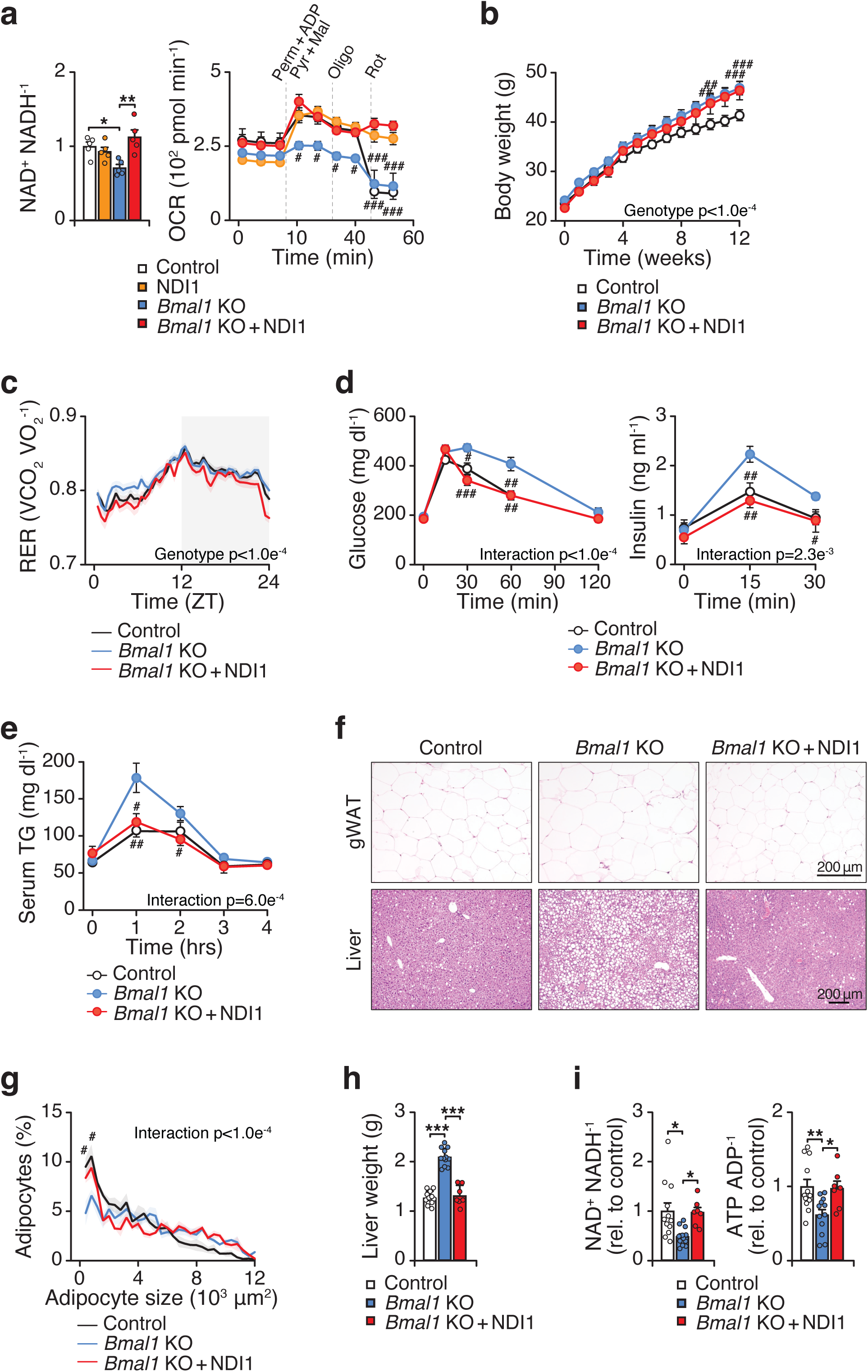
Expression of NDI1 prevents diet-induced metabolic dysfunction in mice lacking the circadian clock in adipocytes. **a**, Intracellular NAD+/NADH ratio and OCR in in the presence of indicated substrates and inhibitors in primary white adipocytes differentiated from Control, NDI1, *Bmal1*-KO, and *Bmal1*-KO + NDI1 mice (n = 5). **b,** Body weight during high fat diet (HFD) feeding for 12 weeks (Control n = 12, *Bmal1*-KO n=12, *Bmal1*-KO + NDI1 n=7). **c,** RER over 24 hours after 10 weeks of HFD feeding (Control n = 12, *Bmal1*-KO n=12, *Bmal1*-KO + NDI1 n=7). **d,** Oral glucose tolerance test (GTT) and insulin during the GTT at 6 weeks of HFD feeding (Control n = 12, *Bmal1*-KO n=12, *Bmal1-*KO + NDI1 n=7). **e,** Oral triglyceride clearance test at 6 weeks of HFD feeding (Control n = 12, *Bmal1*-KO n=12, *Bmal1*-KO + NDI1 n=7). **f,** Representative images of hematoxylin and eosin staining of gWAT and liver after 12 weeks of HFD feeding (10x magnification, scale bars 200 μm). **g,** Distribution in visceral WAT adipocyte size after 12 weeks of HFD feeding (n = 5). **h,** Liver weights after 12 weeks of HFD feeding (Control n = 12, *Bmal1*-KO n=12, *Bmal1*-KO + NDI1 n=7). **i,** The NAD+/NADH and ATP/ADP ratios in WAT after 12 weeks of HFD feeding at ZT14 (Control n = 12, *Bmal1*-KO n=12, *Bmal1*-KO + NDI1 n=7). Data are represented as mean ± SEM. Statistical significance was calculated by one-way ANOVA followed by multiple comparisons in **a** (left panel), **h**, and **i**, two-way ANOVA followed by Dunnett’s multiple comparisons test with *Bmal1*-KO set as the reference group in **a** (right panel), and **b-e** and **g**. The # symbol denotes significance between *Bmal1*-KO and indicated groups. *p < 0.05, **p < 0.01, ***p < 0.001.

During HFD feeding, mice lacking the circadian clock in adipocytes gained excess body weight and had impaired glucose homeostasis, consistent with prior results (**Fig. 4b-e**)^13,16^. Expression of NDI1 in adipocyte-*Bmal1*-KO mice had no effect on weight gain during HFD feeding as compared to adipocyte-*Bmal1*-KO mice (**Fig. 4b**). However, NDI1 expression normalized glucose tolerance, insulin release, and lipid clearance comparable to that of control mice (**Fig. 4d,e**). In NDI1-expressing adipocyte-*Bmal1*-KO mice, we also observed reduced hepatic steatosis and smaller white adipocyte size (**Fig. 4f-h**). Mice lacking *Bmal1* in adipocytes had reduced ratios of NAD+/NADH and ATP/ADP in visceral WAT after HFD feeding, which was restored with NDI1 expression (**Fig. 4i**). This finding is consistent with reduced mitochondrial function in adipocytes during circadian disruption, which is restored with NDI1. Therefore, the impaired metabolic health of adipocyte circadian mutant mice during HFD feeding is alleviated by restoration of mitochondrial NADH dehydrogenase activity through expression of NDI1. Collectively, these findings reveal that circadian disruption in adipocytes promotes metabolic syndrome through disrupting CI function.

To further assess how CI functionality affects transcription during circadian disruption, we analyzed RNA expression in circadian mutant mice with restored CI function. We performed single-nucleus RNA-sequencing in visceral WAT from chow-fed Control, adipocyte-*Bmal1*-KO, and adipocyte-*Bmal1*-KO + NDI1 mice. After doublet removal and filtering of the data, we obtained a total of 12,995 adipocyte nuclei (5,874 Control, 3,949 *Bmal1-*KO, and 3,172 *Bmal1*-KO + NDI1) that grouped into 5 distinct clusters (**Fig. 5a**, **Extended Data Table 2**). The dominant clustering was driven by genotype; however, additional clusters may have been revealed with a greater number of nuclei or with inclusion of other depots, diets, or experimental conditions.

**Figure 5.**
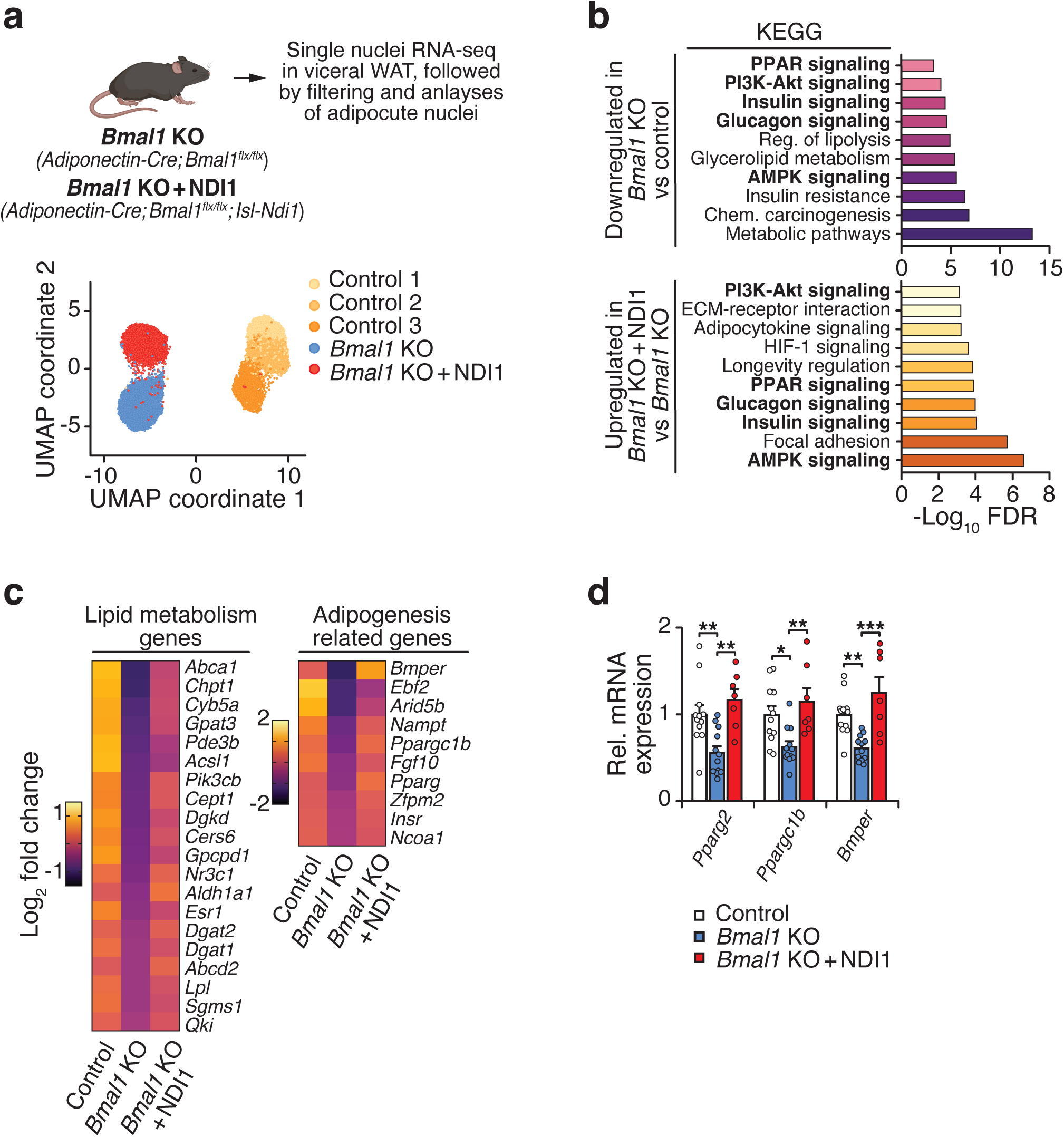
Mitochondrial complex I function regulates the transcriptional landscape in adipocytes. **a**, Single-nuclei RNA-sequencing was performed on visceral WAT from Control (*Bmal1^fx/fx^; NDI1^LSL^*), *Bmal1*-KO (*Adiponectin-Cre; Bmal1^fx/fx^*), and *Bmal1*-KO + NDI1 (*Adiponectin-Cre; Bmal1^fx/fx^; NDI1^LSL^*) mice. Uniform manifold approximation and projection (UMAP) plot showing WAT adipocyte nuclei from Control, *Bmal1*-KO, and *Bmal1*-KO + NDI1 mice (WAT from 4 mice was pooled for each sample). **b,** KEGG pathway analysis of downregulated genes in *Bmal1*-KO vs. Control adipocytes (upper panel) and upregulated genes in *Bmal1*-KO + NDI1 vs. *Bmal1*-KO adipocytes (lower panel). **c,** Heatmap of differentially abundant genes in adipocyte clusters between Control, *Bmal1*-KO, and *Bmal1*-KO + NDI1 mice. **d,** qPCR analysis of indicated genes in visceral WAT from Control, *Bmal1*-KO, and *Bmal1*-KO + NDI1 mice (Control n = 12, *Bmal1*-KO n=12, *Bmal1*-KO + NDI1 n=7). Data are represented as mean ± SEM. Statistical significance was calculated by one-way ANOVA followed by multiple comparisons in **d**. *p < 0.05, **p < 0.01, ***p < 0.001.

Among the adipocyte nuclei in Control WAT, we observed 3 subpopulations with diverse gene expression profiles (**Extended Data Table 3**). Kyoto Encyclopedia of Genes and Genomes (KEGG) pathway analysis revealed that cluster 1 (1,747 nuclei) expressed genes involved in fatty acid metabolism, PI3K-AKT signaling, and lipolysis. Cluster 2 (2,203 nuclei) expressed genes involved in PPAR signaling, insulin signaling, and glycerolipid metabolism. Cluster 3 (1,913 nuclei) was characterized by genes involved in oxidative phosphorylation and reactive oxygen species. Next, we compared gene expression between control and *Bmal1*-KO adipocytes. Adipocytes lacking *Bmal1* had reduced expression of genes involved in fatty acid metabolism, lipid biosynthesis, and fat cell differentiation (**Fig. 5b**). In addition, pathways involved in PPAR, insulin, and glucagon signaling were downregulated in *Bmal1*-KO adipocytes. Notably, *Bmal1*-KO adipocytes had reduced expression of *Pparg*, *Tcf712* (a regulator of lipid and glucose metabolism^33^), and several genes involved in lipid metabolism and adipogenesis (**Extended Data Table 4**, **Fig. 5c**).

Strikingly, pathways involved in metabolism, fat cell differentiation, insulin signaling, and PPAR signaling that were downregulated in *Bmal1*-KO adipocytes were restored with NDI1 expression (**Fig. 5b**). NDI1 expression increased the levels of several adipogenic and metabolic genes, including *Pparg*, *Bmper* (a positive modulator of adipogenesis^34^), *Ppargc1b*, and enzymes involved in lipid metabolism (*Dgat1*, *Dgat2*, *Lpl*) (**Fig. 5c**). qPCR analysis of fractionated adipocytes in lean mice confirmed that NDI1 expression prevents the decline in *Pparg*, *Ppargc1b*, and *Bmper* in mice lacking adipocyte *Bmal1* (**Fig. 5d**). Therefore, restoration of CI function is capable of bypassing some of the major genetic defects involved in lipogenic and nutrient-responsive transcription networks associated with circadian disruption in adipocytes.

## Discussion

In summary, we found that the adipocyte clock programs oxidative metabolism through rhythmic respiration at mitochondrial CI. Dysfunction of CI emerges as a defining feature of circadian disruption and diet-induced obesity in adipose tissue, linking clock misalignment to visceral adipocyte hypertrophy, ectopic lipid deposition in liver, and impaired glucose tolerance. Preserving NADH entry into the respiratory chain with NDI1 prevented these manifestations despite similar weight gain, indicating that mitochondrial electron entry rather than body mass per se constrains metabolic health.

Defects in CI biogenesis are the most common cause of mitochondrial disorders and are linked to age-related diseases, including neurodegeneration^35^. Mechanistically, circadian control of CI likely reflects the intersection of substrate routing and time-of-day coordination of CI biogenesis, stability, and activity. CI is a large 45-subunit L-shaped complex in the mitochondrial inner membrane^19^. It serves as the main gateway for electron entry into the ETC and is the primary site for regeneration of NAD+. Prior work has shown that the clock regulates NAD+ biosynthesis and deacetylation pathways that, in turn, regulate mitochondrial enzyme activity^8,36^. This NAD+ regulation is critical for clock-dependent acetylation, a modification that may influence the assembly and activity of CI^37^. Further work is needed to define the molecular mechanisms through which the clock coordinates CI abundance and function.

In adipose tissue, we observed that CI respiration was rhythmic across gWAT and iWAT, and was reduced by *Bmal1* loss, whereas respiration through CII–CIV was not affected by loss of the clock. The total abundance of CI showed phase- and clock-dependent differences, consistent with temporal regulation of assembly or stability rather than transcriptional control. Because CI is the largest and most energetically costly respiratory complex, coordinating its maintenance with phases of high energetic and cofactor availability may provide an efficient mechanism to sustain rhythmic respiration. In other tissues, including liver, skeletal muscle, and macrophages, loss of *Bmal1* has also been linked to changes in CI, II and III respiration^10,38-41^, highlighting that clock regulation of mitochondrial function is context-dependent and shaped by cell type and metabolic state. In adipocytes, preferential routing of glucose toward lactate production and citrate export for lipogenesis limits succinate flux through CII, which may bias electron entry toward CI^42,43^. We speculate that this reliance on CI-dependent NAD⁺ regeneration, together with the dynamic demands of lipid synthesis and storage, makes CI sensitive to circadian regulation in adipose tissue.

Mitochondria have emerged as dynamic signaling organelles that regulate cellular function beyond their metabolic roles^44^. Loss of CI resulted in reduced expression of transcription networks involved in insulin sensitivity, PPAR signaling, and fatty acid metabolism. Both genetic circadian disruption and HFD feeding shifted adipocyte cell state from a pro-adipogenic and lipogenic gene signature toward an anti-adipogenic and insulin resistant state^45,46^. Preserving CI function with NDI1 expression maintained adipogenic and lipogenic transcription in adipocytes lacking the circadian clock. Therefore, mitochondrial dysfunction mediates many of the gene expression defects caused by circadian disruption. NDI1 expression led to increased NAD+ and α-ketoglutarate with reduced succinate. Therefore, the retrograde signal may involve altered activity of chromatin-modifying enzymes sensitive to these metabolites, such as NAD+-dependent deacetylases and α-ketoglutarate-dependent demethylases. Defining the mitochondrial signals linking CI respiration to transcription will be key to understanding how circadian clocks coordinate cellular identity with metabolic state. All of our experiments were performed in male mice; given known sex differences in metabolic and circadian physiology, whether female mice exhibit similar regulation of CI in adipose tissue remains to be determined.

Collectively, our studies identify an essential function of the circadian clock in mitochondrial respiration through CI. These findings extend both insight into clock transcriptional mechanisms in adipose bioenergetics, and more broadly, into how repair of mitochondrial metabolism is sufficient to reprogram transcription.

## Methods

### Animals and diets

All animal experiments were performed according to procedures approved by the Northwestern University Institutional Animal Care and Use Committee. All experiments were performed using male mice on a C57BL/6J background. *Adiponectin-Cre* mice (Jackson Laboratories #028020) were bred to *Bmal1*^fx/fx^ (Dr. C. Bradfield, University of Wisconsin), *NDI1^LSL^* (Dr. N. Chandel, Northwestern University)^25^, and *Ndufs2^fx/fx^* (Dr. J. López-Barneo, Universidad de Sevilla) mice. mPer2^Luc^ (Jackson Laboratories # 006852) mice were used for in vitro circadian synchronization experiments. Mice were maintained at room temperature (23-25°C) on a standard chow diet with a 12 -hour light/dark cycle and free access to water and food unless specified. In HFD feeding experiments, mice were fed HFD (45% kcal from fat, Bio-Serv) beginning at 3 months of age.

### Indirect calorimetry

Mice were singly housed in Sable Systems metabolic cages at room temperature (24°C). Mice were acclimated to the metabolic cages for 5 days and were provided ad libitum access to food and water. Food intake, VO_2_, VCO_2_, RER (VCO_2_/VO_2_), and locomotor activity (beam breaks) were continuously monitored throughout the experiment.

### Metabolic phenotyping

For glucose tolerance tests, mice were fasted for 2 hours (beginning at ZT0) and then administered glucose by oral gavage (2.5 g/kg body weight). At the indicated time-points, tail blood was collected. Blood glucose was measured using a Bayer Contour glucometer. Serum insulin was measured using Ultra-Sensitive Mouse Insulin ELISA (Crystal Chem). For triglyceride clearance tests, mice were fasted overnight (∼15 hours), then administered 20% intralipid (Sigma) by oral gavage (15 µl/g body weight) at ZT2. Blood was collected hourly and then assayed for serum triglycerides (Infinity Triglycerides Reagent, Thermo).

### Histological analysis

Adipose tissues and liver were harvested and placed in 4% paraformaldehyde overnight, followed by 70% ethanol for up to 4 days before processing. Paraffin embedding, tissue sectioning, and hematoxylin and eosin (H&E) staining were performed by the Mouse Histology and Phenotyping Laboratory core at Northwestern University. Images were obtained on a Keyence BZ-X810. Adipocyte size analysis was performed using the Keyence BZ-X Analyzer software based on bright-field images of H&E-stained paraffin sections.

### Gene expression analysis by quantitative PCR

RNA was isolated from frozen tissues or fractionated adipocytes using Trizol and the RNeasy mini kit (Qiagen). To isolate fractionated adipocytes, fresh visceral WAT was minced and digested for 1 hour in PBS containing 10 mM CaCl2, 2.4 units/mL Dispase II (Roche), and 1.5 units/mL collagenase D (Roche). The homogenate was filtered through a 100 μm cell strainer and centrifuged at 500 xg, followed by transfer of the floated adipocytes to an Eppendorf tube for RNA isolation. cDNA was synthesized using M-MLV Reverse Transcriptase (Invitrogen) and Random Primers (Invitrogen). Relative mRNA levels were determined by quantitative PCR using iTaq Universal SYBR® Green Supermix (Bio-Rad). Values were normalized to three housekeeping genes (*Actb, Rplp0, and Ppia*) using the ΔΔ-Ct method. All primer sequences are listed within **Extended Data Table 5**.

### WAT mitochondria isolation

To isolate mitochondria, fresh visceral WAT was minced then homogenized using a drill-operated Teflon pestle in ice-cold MSHE buffer (70 mM sucrose, 210 mM mannitol, 5 mM HEPES, 1 mM EDTA, pH 7.2) containing 0.5% FA-free BSA. The homogenate was centrifuged at 800 xg for 10 min, followed by centrifugation of the supernatant at 8,000 xg for 10 min. The mitochondrial pellet was resuspended in MSHE buffer with BSA and protein concentration was determined using Piece BCA Protein Assay Kit (Thermo).

### Mitochondrial function and respiration

Isolated WAT mitochondria, cultured primary differentiated adipocytes, and MEFs were assayed for oxygen consumption rate (OCR) using a Seahorse XFe96 Extracellular Flux Analyzer (Agilent). For mitochondrial OCR assays, mitochondria were plated on Seahorse Biosciences 96-well culture plates (5 μg of protein per well) and centrifuged for 20 min at 2000 xg at 4°C to adhere mitochondria to base of wells. For electron coupling assays, respiratory substrates (10 mM pyruvate + 2 mM malate, 10 mM succinate + 2 μM rotenone, 80 μM palmitoyl- or octanoyl-carnitine + 0.5 mM malate, or 10 mM glutamate + 10 mM malate) were diluted in MAS buffer with BSA (70 mM sucrose, 220 mM mannitol, 10 mM KH_2_PO_4_, 5 mM MgCl_2_, 2 mM HEPES, and 1 mM EGTA, pH 7.2 in 0.2% FA-free BSA) and added to mitochondria. Substrate injection was as follows: 4 mM ADP at port A, 10 μM oligomycin at port B, 10 μM carbonyl cyanide 4-(trifluoromethoxy)phenylhydrazone (FCCP) at port C, and 10 μM antimycin A at port D. For determination of CI activity, MAS plus alamethicin (2.5 μg/ml) and cytochrome C (10 μg/ml) were added onto the mitochondria followed by: 5 mM pyruvate + 5 mM malate + 1mM NADH at port A, 2 μM rotenone + 4 μM antimycin A at port B, 0.5 mM TMPD + 1 mM ascorbic acid at port C, and 50 mM azide at port D. For electron flow assays, mitochondria were plated in MAS buffer with 10 mM pyruvate + 2 mM malate + 4 μM FCCP followed by: 20 μM rotenone in port A, 100 mM succinate in port B, 40 μM antimycin A in port C, and 1 mM TMPD + 100 mM ascorbic acid in port D.

For OCR assays in cells, culture media was replaced with MAS buffer with 4% BSA, pH 7.2. Activity measurements for respiratory chain CI-IV in permeabilized cells were performed as previously described^47^. The first injection in port A contained the substrate for each individual respiratory chain complex (I: 5 mM pyruvate + 2.5 mM malate, II: 10 mM succinate + 1 μM rotenone, III: 0.5 mM duroquinol, IV: 0.5 mM TMPD + 2 mM ascorbic acid) along with 1 mM ADP and 10 nM recombinant perfringolysin O (Agilent). For determination of CI activity (NADH-dependent OCR), MAS plus alamethicin (2.5 μg/ml) and cytochrome C (10 μg/ml) was added onto the cells followed by: 5 mM pyruvate + 5 mM malate + 1mM NADH + 10 nM recombinant perfringolysin O at port A, 2 μM rotenone + 4 μM antimycin A at port B, 0.5 mM TMPD + 1 mM ascorbic acid at port C, and 50 mM azide at port D. OCR of MEFs was normalized to total protein content due to differences in proliferation.

### Western blotting

Protein extracts from adipose tissue mitochondria were prepared by homogenization in RIPA buffer containing protease inhibitors (Sigma). The homogenate was spun at 18,000 xg for 15 minutes and the supernatant was used for subsequent analysis. Protein levels were quantified using BCA protein assay. Protein extracts were then subject to SDS-PAGE gel electrophoresis and transferred to Immobilin-FL PVDF transfer membrane. The blots were incubated with primary antibody at 4°C overnight followed by incubation with IR Dye-coupled secondary antibody and visualization by the LI-COR Odyssey Fc imaging system.

### BN-PAGE

For BN-PAGE, isolated mitochondria were resuspended in buffer (20 mM Tris pH 7.4; 0.1 mM EDTA; 50 mM NaCl; 10% [v/v] glycerol) containing 2% n-dodecyl b-D-maltoside (DDM) for a 20-minute incubation on ice, followed by centrifugation at 20,000 xg for 10 minutes. Protein concentration of the supernatant was measured using BCA protein assay. 15 μg of protein was mixed with NativePAGE sample buffer (Invitrogen), Coomassie G-250, and protease inhibitor, loaded into lanes of a 3-12% Bis-Tris NativePAGE gel (Invitrogen), and run according to the manufacturer’s instructions. Proteins were visualized using western blot analysis.

### Mouse Embryonic Fibroblast (MEF) cell culture

Control, *Bmal1^−/−^*, *Cry1^−/−^;Cry2^−/−^* MEFs were cultured in Dulbecco’s Modified Eagle’s Medium (DMEM) (Sigma) supplemented with 15% fetal bovine serum (Atlanta Biological) and 1% penicillin-streptomycin at 37°C with 5% CO2. Culture medium was exchanged every one to two days. All cells used in these experiments were <15 passages.

### Isolation of WAT stromal vascular cells and adipocyte differentiation

To isolate the stromal vascular fraction (SVF), minced inguinal WAT (combined depots from 4-5 mice per genotype) from 4-6 week old mice was digested in 1 mg/mL collagenase D with 1.5% BSA in sterile Hank’s Balanced Salt Solution (HBSS) and incubated in a 37°C shaking water bath for 1 hr. The mixture was then passed through a 100 µm cell strainer, pelleted at 500 xg, resuspended in PBS, passed through a 40 µm cell strainer, and again pelleted at 500 xg. SVF cells were expanded in growth media containing DMEM/F12 with GlutaMAX (Thermo) and 10% FBS with 100 U/mL Pen/Strep, and 50 μg/ml gentamicin, and incubated in 10% CO2 at 37°C. Upon reaching confluence, cultures were incubated with the adipogenesis induction cocktail (growth media supplemented with 5 μg/ml insulin, 1 μM dexamethasone, 0.5 mM isobutylmethyxanthine, and 1 μM rosiglitazone) for day 1-2 of differentiation. For days 2-4 of differentiation, the cells were incubated with 5 µg/ml insulin and 1 µM rosiglitazone. For days 4-8 of differentiation, the cells were maintained in growth media supplemented with 5 µg/ml insulin until harvest. Adipocytes were considered mature for use in assays on day 8 of differentiation. For synchronization experiments, cells were synchronized with dexamethasone (100 nM, 30 min) on day 6 of differentiation and assayed or collected at the indicated timepoints. In parallel, Per2-luciferase rhythms were monitored in synchronized cells cultured in DMEM containing 0.1 mM luciferin using a LumiCycle apparatus (Actimetrics).

### Isolation of polar metabolites

For metabolomics of tissues, 100-200 mg of weighed tissue was homogenized in 1 mL of ice-cold 80% methanol with a Qiagen Tissue Lyser II (3 sets of 2 minutes on maximum speed, with 2 minutes of rest on dry ice in between sets). The homogenized tissue was then incubated at -80°C for 15 minutes and then centrifuged at 18,000 xg at 4°C for 5 minutes. The lipid layer was avoided, and the supernatant was transferred to a new tube on dry ice and then centrifuged at 18,000 xg at 4°C for 5 minutes. The equivalent volume of 20 mg of tissue was transferred to a new tube, which was then dried in a speed-vac.

For metabolomics of cell cultures, cells were differentiated in 6-well plates. On day 6 of differentiation, media was changed to glucose/phenol red/glutamine/pyruvate-free DMEM (Thermo) supplemented with 0.5% FBS, 4.5 g/L glucose, 4 mM glutamine, 0.5 mM pyruvate, and 5 μg/ml insulin. After 48 hours, the media was removed, 400 µL of ice-cold 80% methanol was added, and the cells were incubated at -80°C for 15 minutes. Lysed cells were scraped on dry ice and transferred to Eppendorf tubes for centrifugation at 18,000 xg at 4°C for 5 minutes. The entire supernatant was transferred to a new tube and dried in a speed-vac. The pellet was resuspended in 200 µL of 8M Urea with 10 mM tris (pH=8), incubated at 60C for 30 min with shaking, and centrifuged for 15 minutes at 18,000 xg for protein quantification of the supernatant with DC Protein Assay (Bio-Rad).

### LCMS analysis of metabolites

The dried metabolites were reconstituted in 60% acetonitrile followed by overtaxing for 30 seconds and then centrifugation for 30 min at 20,000 xg at 4 °C. Supernatant was analyzed by High-Performance Liquid Chromatography and High-Resolution Mass Spectrometry and Tandem Mass Spectrometry (HPLC-MS/MS). Specifically, system consisted of a Thermo Q-Exactive in line with an electrospray source and an Ultimate3000 series HPLC consisting of a binary pump, degasser, and auto-sampler outfitted with a Xbridge Amide column (Waters; dimensions of 3.0 mm × 100 mm and a 3.5 µm particle size). The mobile phase A contained 95% (vol/vol) water, 5% (vol/vol) acetonitrile, 10 mM ammonium hydroxide, 10 mM ammonium acetate, pH = 9.0; B was 100% Acetonitrile. The gradient was as following: 0 min, 15% A; 2.5 min, 30% A; 7 min, 43% A; 16 min, 62% A; 16.1-18 min, 75% A; 18-25 min, 15% A with a flow rate of 150 μL/min. The capillary of the ESI source was set to 275 °C, with sheath gas at 35 arbitrary units, auxiliary gas at 5 arbitrary units and the spray voltage at 4.0 kV. In positive/negative polarity switching mode, an *m*/*z* scan range from 60 to 900 was chosen and MS1 data was collected at a resolution of 70,000. The automatic gain control (AGC) target was set at 1 × 10^6^ and the maximum injection time was 200 ms. The top 5 precursor ions were subsequently fragmented, in a data-dependent manner, using the higher energy collisional dissociation (HCD) cell set to 30% normalized collision energy in MS2 at a resolution power of 17,500. Besides matching m/z, metabolites are identified by matching either retention time with analytical standards and/or MS2 fragmentation pattern. Data acquisition and analysis were carried out by Xcalibur 4.1 software and Tracefinder 4.1 software, respectively. Metabolites were normalized to total ion count of each sample. NAD+/NADH and ATP/ADP ratios were calculated from the peak area values of each metabolite within the same individual sample and compared between groups.

### RNA-sequencing from fractionated adipocytes, library preparation, and analysis

RNA was isolated from fractionated adipocytes as described above. RNA quality was assessed by Tapestation and those with an RNA Integrity Number (RIN) greater than 8 were used for library prep. Libraries were constructed from 250 ng of RNA using the NEBNext RNA Ultra II Directional library prep kit (NEB). Average library size and concentration were determined by Tapestation and qPCR (NEBNext Library Quant kit, NEB), respectively, prior to pooling. Pooled libraries were sequenced on a NextSeq 2000 (Illumina) using 100 bp single-end sequencing. Libraries were sequenced to an average depth of ∼20M aligned reads for differential gene analyses. Sequences were aligned to the mm39 transcriptome with STAR using the –quantMode Transcriptome and GeneCounts options. Gene counts were obtained from mm39 annotations (Ensembl 112) using rsem-calculate-expression. Differential expression was carried out using DESeq2 with default parameters, and pathway analysis was performed using pathfindR.

### Single-nuclei RNA-sequencing and analysis

For isolation of nuclei from visceral WAT, fresh WAT was minced in nucleus preparation buffer (NPB; 10 mM HEPES, 1.5 mM MgCl_2_, 10 mM KCl, 250 mM sucrose, 0.001% NP-40) with 0.2 U/ul Roche Protector RNase inhibitor followed by dounce homogenization. The homogenate was filtered through a 100 μM cell strainer and centrifuged at 1000 xg for 10 minutes at 4°C, then the nuclear pellet was washed once in NPB followed by resuspension in PBS with 1% BSA and 0.2 U/ul Roche Protector RNase inhibitor. Single-nuclei RNA-seq libraries were prepared using 10X Genomics Chromium Next GEM Single Cell 3’ Reagent Kits v.3.1 with loading of ∼12,000 total nuclei per sample and sequenced on the Illumina HiSeq 4000 instrument.

Cell Ranger software was used to perform demultiplexing and align reads to mm39. Nuclei were filtered using Seurat based on the total number of genes and molecules, as well as percent of mitochondrial reads (500 < nFeature_RNA < 6000; 1000 < nCount_RNA < 25000; percent.mt < 15%). Dimensionality reduction was performed in Seurat using Uniform Manifold Approximation and Projection (UMAP). Differential gene expression between clusters was performed using FindMarkers function. Adipocyte nuclei were identified based on the expression of established markers (*Lipe*, *Plin4*, *Plin1*, and *Pparg*) and filtered for re-clustering using Seurat. UMAP plots of the adipocyte nuclei were visualized using Loupe Browser of the imported analysis from Seurat. KEGG pathway enrichment analysis of differentially expressed genes between clusters or samples was performed using limma and biomaRt.

### Statistical Analyses

Data was analyzed using R (v4.3.2) for sequencing data and Prism (v9.0) for statistical significance all other data. Results are presented as mean ± standard error of the mean (S.E.M.) for all replicates included in each analysis. Statistical tests and significance thresholds are specified in figure legends.

## Data Availability

Data in this study will be made publicly available by publication date in NCBI’s Gene Expression Omnibus (GEO) and are accessible through the GEO Series accession number GSE245850.

## Acknowledgements

We thank the following core facilities at Northwestern University: Robert H. Lurie Cancer Center (RHLCCC) Metabolomics Core, Mouse Histology and Phenotyping Laboratory (MHPL), and NUSeq Core Facility. We would like to acknowledge NIH grant 1S10OD025120 for the 10x Chromium housed in the NUSeq Facility, and NIH grant NCI P30-CA060553 awarded to the RHLCCC which supports the MHPL. We thank all members of the Bass, Barish, Beutler, and Peek laboratories for helpful discussions, P. Gao for performing metabolomics, and C. Wai for preparing the 10X genomics libraries.

Research support was from the NIH National Institute of Diabetes and Digestive and Kidney Diseases (NIDDK) grants R01DK127800, R01DK113011, R01DK090625, R01DK132647, and P30DK020595 and National Institute on Aging (NIA) grants R01AG065988 and P01AG011412 to J.B.; NIDDK grant F32DK122675 and American Diabetes Association Pathway to Stop Diabetes award 1-24-INI-01 to C.H.; NIDDK grant F30DK116481 (to B.J.W.), NIDDK grant F31DK130589 to N.J.W., National Heart, Lung, and Blood Institute (NHLBI) T32HL076139 to C.R.R., NIDDK grant R01DK123358 to C.B.P., NIDDK grant R01DK108987 and Veteran’s Affairs grant I01BX004898 to G.D.B., and National Cancer Institute (NCI) grant R35CA197532, NHLBI grant P01HL071643, and NIA grant P01AG049665 to N.S.C.

## Author contributions

C.H. and J.B. conceived the study and wrote the manuscript. C.H., N.W., B.J.W., Y.C., J.H., Z.Z., C.B.P, G.D.B., N.S.C., and J.B. designed experiments and interpreted data. C.H., N.J.W., B.J.W., Y.C., J.H., R.N., and J.V.M. performed experiments and analyzed data. B.M. visualized the data. C.R.R. generated the *NDI1^LS^*^L^ mice. C.H., N.W., B.J.W., Y.C., J.H., Z.Z., A.K.T., C.R.R., J.C., K.M.R., C.B.P, G.D.B., N.S.C., and J.B edited the manuscript.

---

**Extended Data Figure 1.**
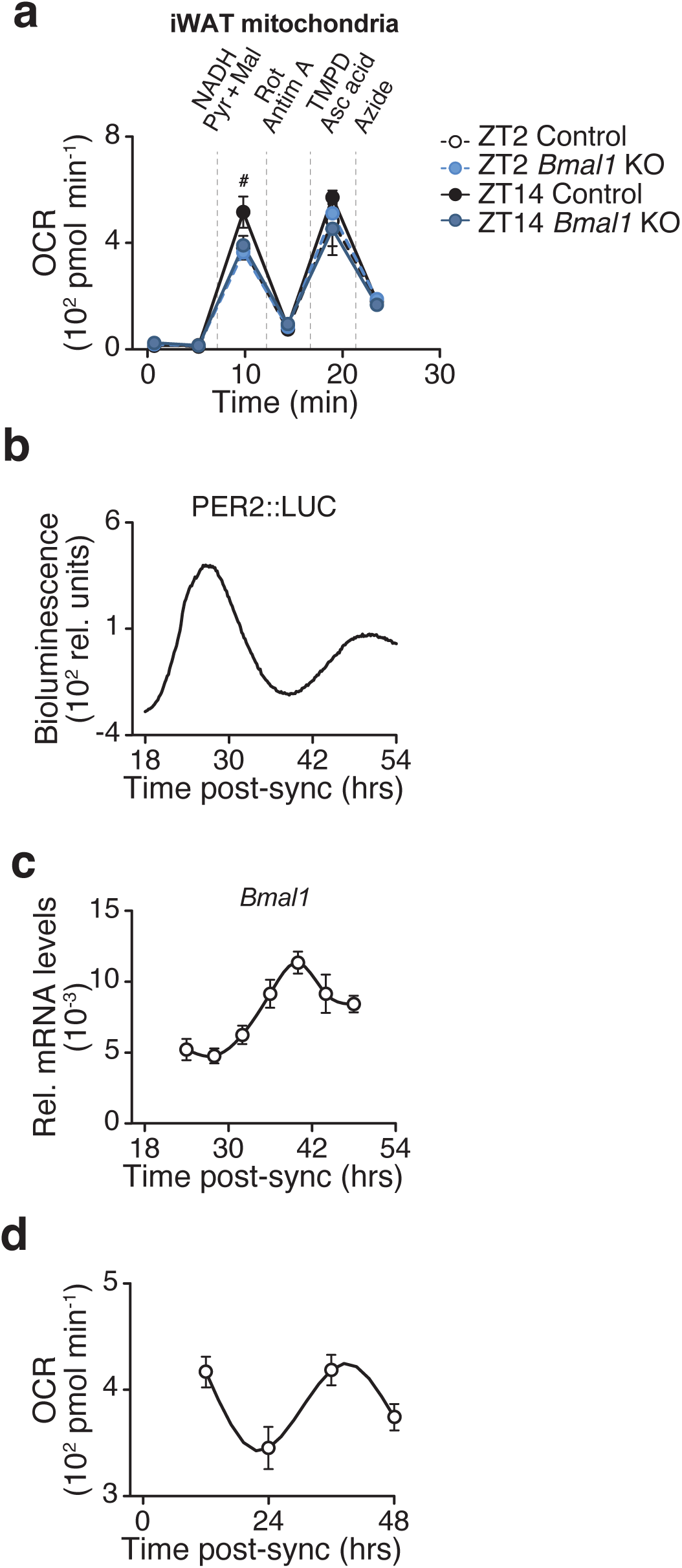
The circadian clock regulates the abundance of mitochondrial complex I, related to Figure 1. **a,** OCR in iWAT mitochondria of Control and *Bmal1*-KO mice at ZT2 and ZT14 in the presence of indicated substrates and inhibitors (n = 5). TMPD, Tetramethyl-p-phenylenediamine; Asc acid; ascorbic acid. **b,** Representative bioluminescence of PER2::LUC oscillations in synchronized primary white adipocytes from mPer2:Luc mice. **c,** Relative mRNA expression of Bmal1 in synchronized white adipocytes in vitro (n = 6). **d,** NADH-dependent respiration in synchronized white adipocytes in vitro (n=6). Data are represented as mean ± SEM. Statistical significance was calculated by two-way ANOVA followed by Dunnett’s multiple comparisons test with ZT14 Control set as the reference group in **a**.

**Extended Data Figure 2.**
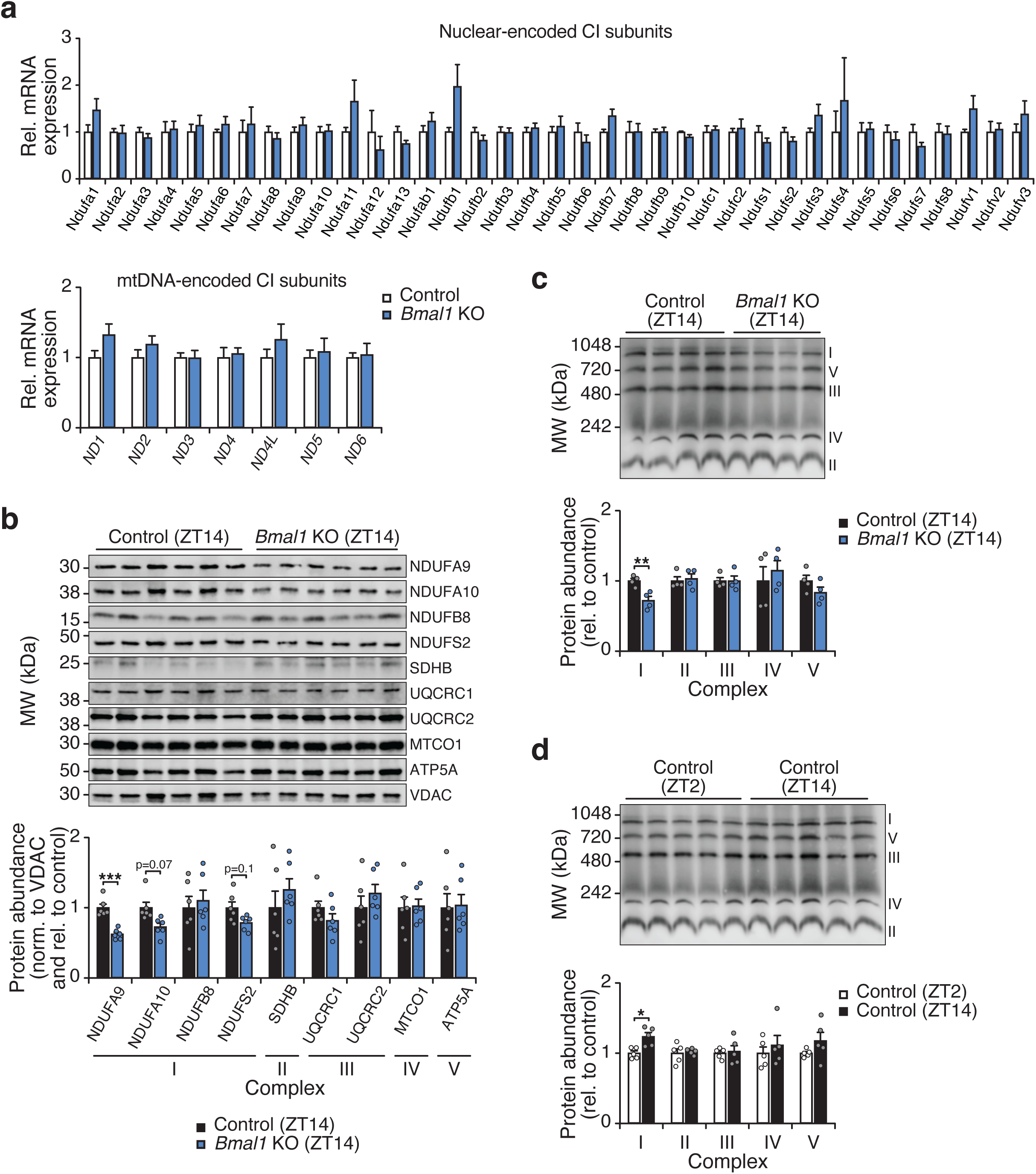
The circadian clock regulates the abundance of mitochondrial complex I, related to Figure 1. **a,** mRNA abundance of complex I subunits in adipocytes from the visceral WAT depot of Control (*Bmal1^fx/fx^*) and *Bmal1*-KO (*Adiponectin-Cre; Bmal1^fx/fx^*) mice (n=6). **b,** SDS-PAGE immunoblot of electron transport chain proteins in isolated WAT mitochondria from Control and *Bmal1-KO* mice harvested at the indicated timepoints (n=6). **c,** BN-PAGE immunoblot on isolated WAT mitochondria from Control and *Bmal1-KO* mice at ZT14 using an antibody mix against OXPHOS complex subunits (n=4). **d,** BN-PAGE immunoblot on isolated WAT mitochondria from Control mice at ZT2 and ZT14 using an antibody mix against OXPHOS complex subunits (n=5). Data are represented as mean ± SEM. Statistical significance was calculated by two-way ANOVA followed by multiple comparisons.

**Extended Data Figure 3.**
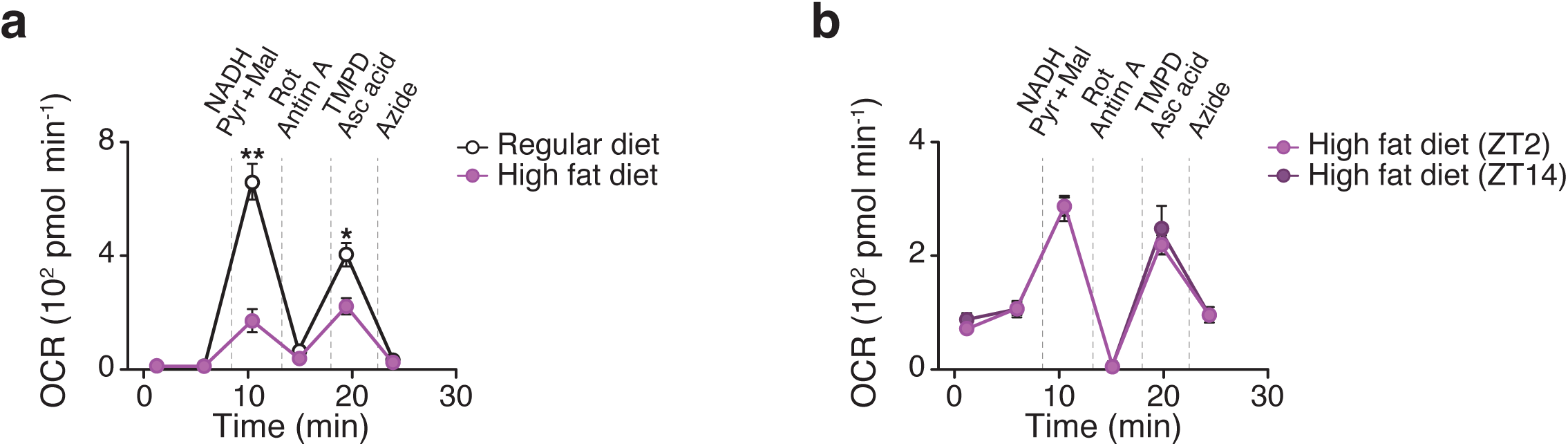
High fat diet feeding leads to reduced and loss of diurnal CI respiration in visceral WAT mitochondria, related to Figure 2. **a,** OCR in WAT mitochondria of wildtype mice at ZT14 following 12 weeks of chow or HFD feeding. Mitochondria were treated with pyruvate/malate/FCCP followed sequentially with rotenone, succinate, antimycin A, and N,N,N′,N′-Tetramethyl-p-phenylenediamine (TMPD)/ascorbic acid (n = 5). **b,** OCR in WAT mitochondria of wildtype mice at ZT2 and ZT14 following 12 weeks of HFD feeding. Mitochondria were treated with pyruvate/malate/NADH followed sequentially with rotenone/antimycin A, TMPD/ascorbic acid, and azide (n = 6). Data are represented as mean ± SEM. Statistical significance was calculated by two-way ANOVA followed by multiple comparisons in **a-b**. **p < 0.01.

**Extended Data Figure 4.**
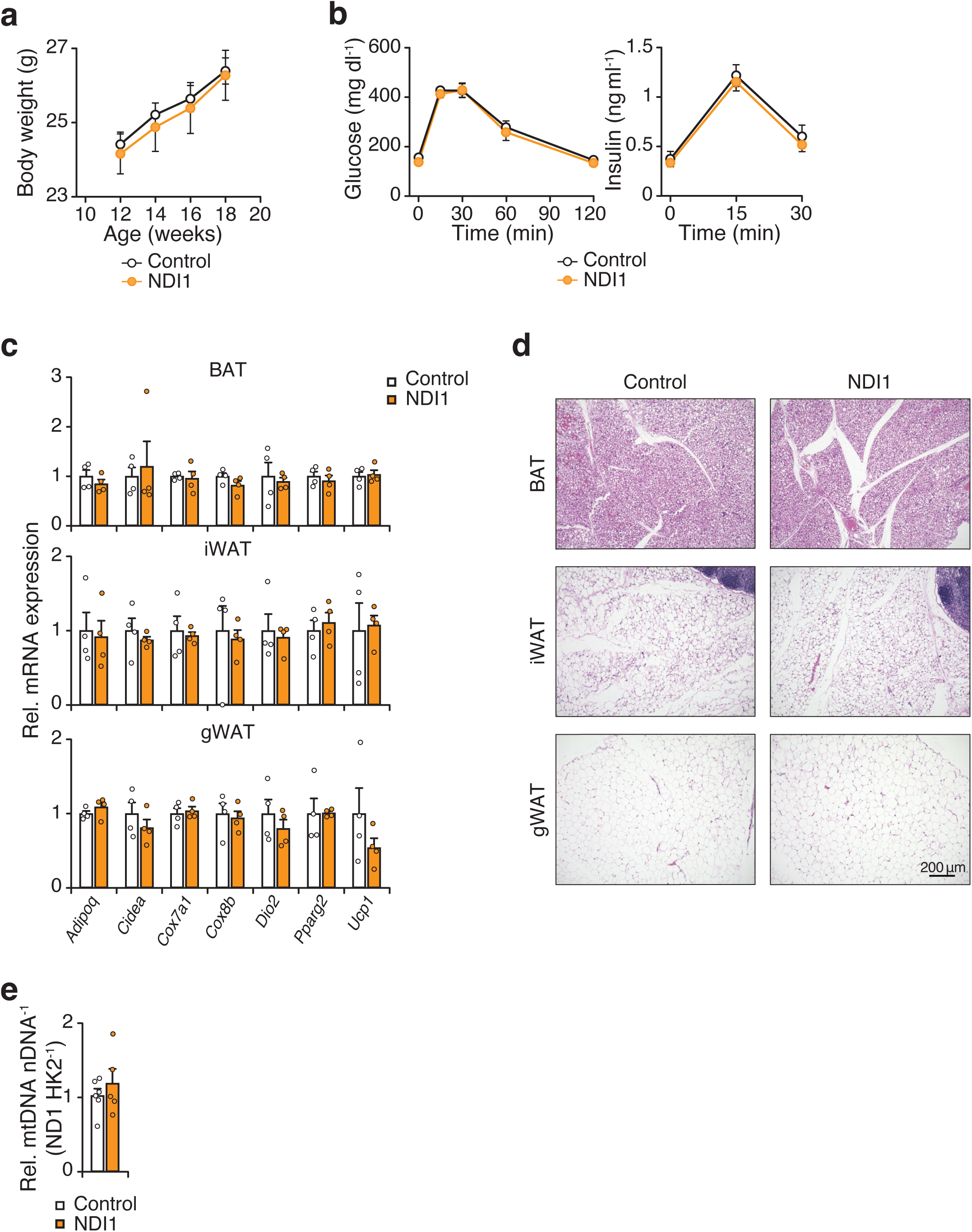
NDI1 expression does not affect adipocyte physiology or glucose homeostasis in chow-fed mice, related to Figure 2. **a,** Body weight of Control (*NDI1^LSL^*) and NDI1 (*Adiponectin-Cre; NDI1^LSL^*) mice at their indicated age on a chow diet (n = 5-6). **b,** Glucose tolerance test (GTT) and insulin during the GTT at 3 months of age (n = 5-6). **c,** Gene expression of indicated genes involved in thermogenesis in brown adipose tissue (BAT), inguinal white adipose tissue (iWAT), and gonadal white adipose tissue (gWAT) of Control and NDI1 mice at 3 months of age (n = 4). **d,** Representative images of hematoxylin and eosin staining of indicated adipose tissue depots in Control and NDI1 mice at 3 months of age (10x magnification, scale bars 200 μm). **e,** Mitochondria/nuclear DNA ratio in gWAT from control and NDI1 mice at 3 months of age (n = 5-6). Data are represented as mean ± SEM. Statistical significance was calculated by two-way ANOVA followed by multiple comparisons in **a-c** and unpaired t-test in **e**.

**Extended Data Figure 5.**
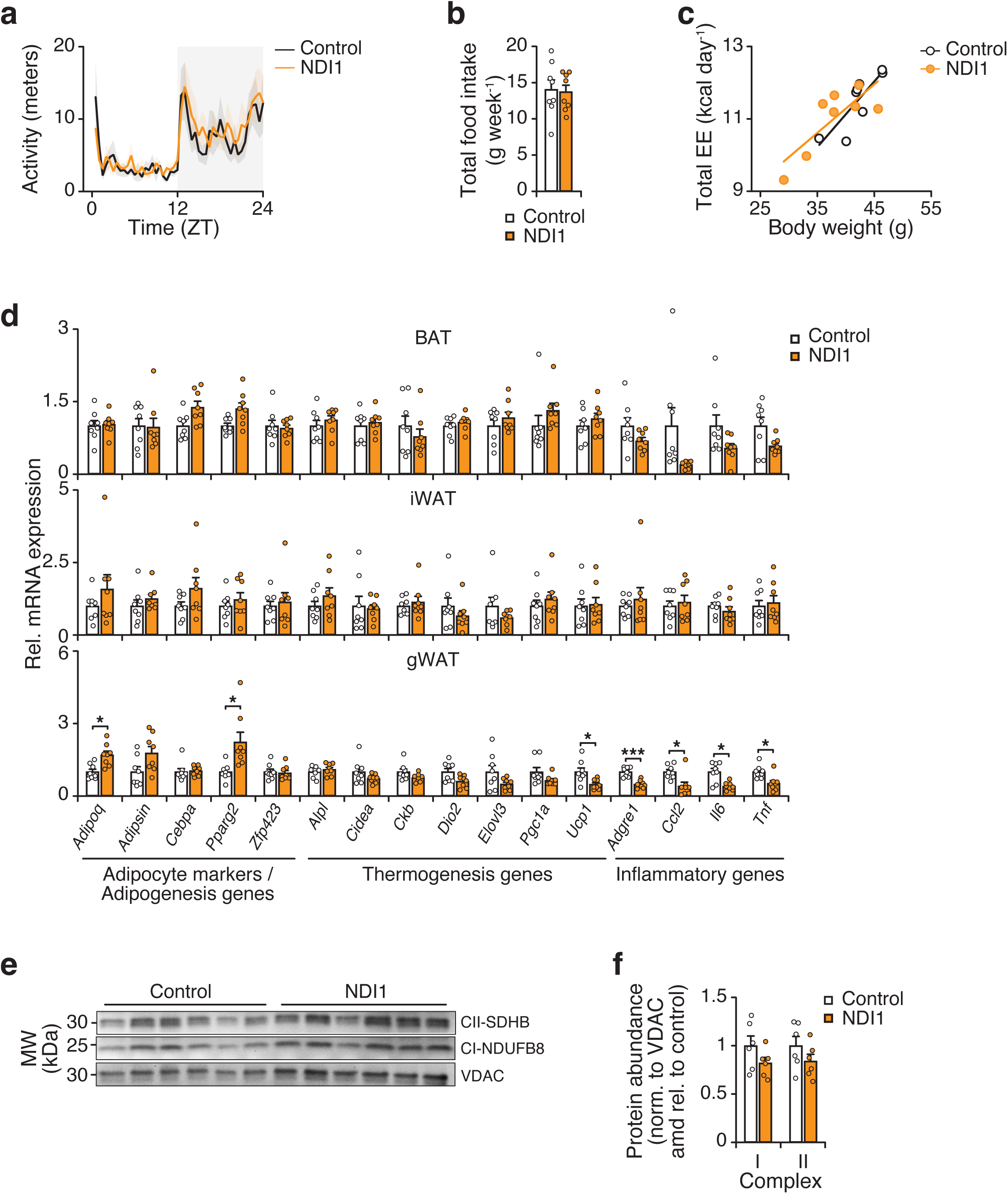
Expression of NDI1 does not affect activity, food intake, or energy expenditure during high fat diet feeding, related to Figure 2. **a,** Daily activity rhythms of Control and NDI1 mice after 10 weeks of HFD feeding (n = 8). **b,** Total daily food intake in Control and NDI1 mice after 10 weeks of HFD feeding (n = 8). **c,** Total energy expenditure over 24 hours versus body weight after 10 weeks of HFD feeding (n = 8). **d,** Gene expression of indicated genes involved in thermogenesis in BAT, inguinal WAT, and gonadal WAT from Control and NDI1 mice after 12 weeks of HFD feeding (n = 8). **e,** SDS-PAGE immunoblot of complex I and II subunits in isolated gWAT mitochondria from Control and NDI1 mice after 10 weeks of HFD feeding (n=6). **f,** Mitochondria/nuclear DNA ratio in gWAT from control and NDI1 mice after 10 weeks of HFD feeding (n = 6). Data are represented as mean ± SEM. Statistical significance was calculated by two-way ANOVA followed by multiple comparisons in **a**, **d**, and **f**, unpaired t-test in **b** and ANCOVA in **c**. *p < 0.05, ***p < 0.001.

**Extended Data Figure 6.**
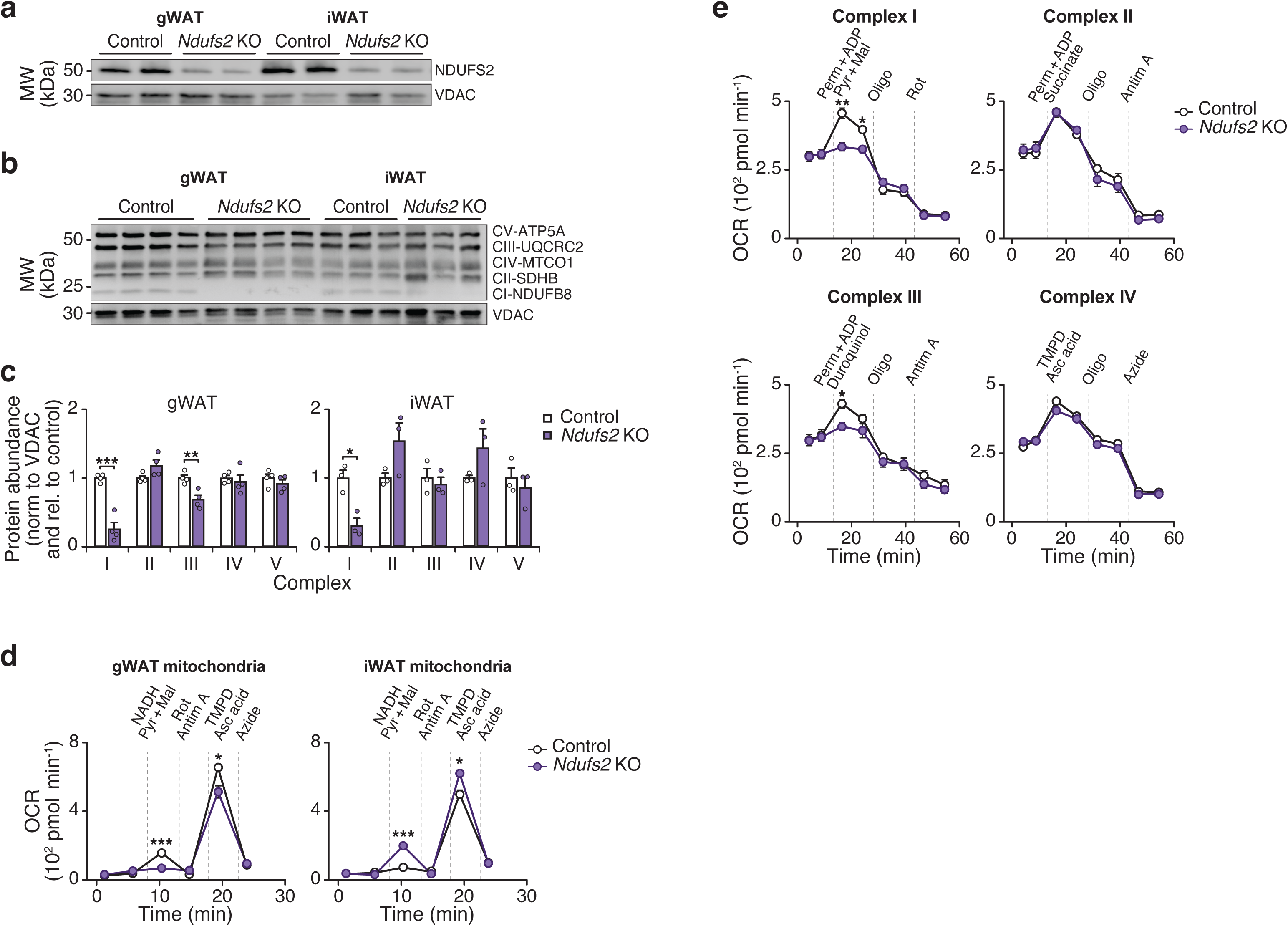
Loss of *Ndufs2* in adipocytes leads to reduced complex I and III activity in gWAT, related to Figure 3. **a,** SDS-PAGE immunoblot of *Ndufs2* in isolated mitochondria from Control (*Ndufs2^fx/fx^*) and *Ndufs2*-KO (*Adiponectin-Cre; Ndufs2^fx/fx^*) mice at 3 months of age on a chow diet (n = 2). **b,** SDS-PAGE immunoblot of electron transport chain proteins in isolated gWAT mitochondria from Control and *Ndufs2*-KO mice at 3 months of age (n = 3-4). **c,** Quantification of protein abundance of samples in B. **d,** OCR in gWAT mitochondria of Control and *Ndufs2*-KO mice at ZT14 in the presence of indicated substrates and inhibitors (n = 7-7). TMPD, Tetramethyl-p-phenylenediamine; Asc acid; ascorbic acid. **e,** OCR of differentiated adipocytes from Control and *Ndufs2*-KO mice in the presence of permeabilizer (PERM), ADP, and complex I to IV substrates followed sequentially by oligomycin and antimycin A (n = 5). Data are represented as mean ± SEM. Statistical significance was calculated by two-way ANOVA followed by multiple comparisons in **c**, **d**, and **e**.*p < 0.05, **p < 0.01, ***p < 0.001.

**Extended Data Figure 7.**
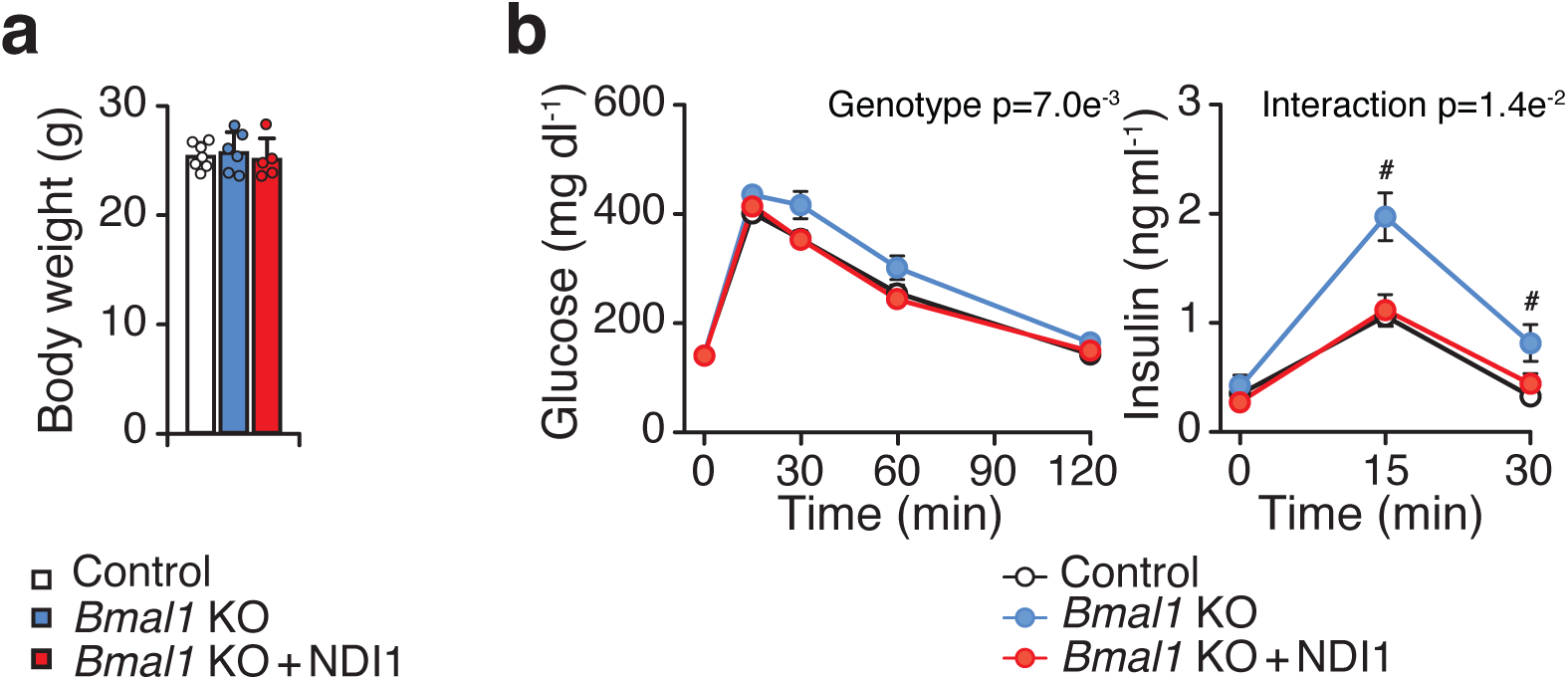
Deletion of Bmal1 leads to impaired glucose tolerance on a chow diet, which is prevented by NDI1, related to Figure 4. **a,** Body weight of Control, Bmal1-KO, and Bmal1-KO + NDI1 mice at 4 months of age on a chow diet (n = 5-7). (B) Glucose tolerance test (GTT) and insulin during the GTT of the mice in A at 4 months of age (n = 5-7). Data are represented as mean ± SEM. Statistical significance was calculated by one-way ANOVA in **a** and two-way ANOVA followed by Dunnett’s multiple comparisons test with Bmal1-KO mice set as the reference group in **b**. # denotes significance between Bmal1-KO vs other groups (p < 0.05).

## Tables

**Extended Data Table 1.** RNA-sequencing analysis of Ndusf2-KO and Bmal1-KO fractionated adipocytes, related to Figure 3

**Extended Data Table 2.** Top 50 differentially expressed genes in adipocyte clusters, related to Figure 3

**Extended Data Table 3.** Gene and pathway lists for adipocyte clusters in control mice, related to Figure 3

**Extended Data Table 4.** Differentially expressed genes between samples, Related to Figure 3

**Extended Data Table 5.** Primer sequences for qPCR, related to STAR Methods

## References

1 Laothamatas, I., Rasmussen, E. S., Green, C. B. & Takahashi, J. S. Metabolic and chemical architecture of the mammalian circadian clock. Cell Chem Biol 30, 1033–1052 (2023). 10.1016/j.chembiol.2023.08.014

2 Wang, F. et al. Meta-analysis on night shift work and risk of metabolic syndrome. Obes Rev 15, 709–720 (2014). 10.1111/obr.12194

3 Turek, F. W. et al. Obesity and metabolic syndrome in circadian Clock mutant mice. Science 308, 1043–1045 (2005). https://doi.org:1108750 [pii] 10.1126/science.1108750

4 Pariollaud, M. & Lamia, K. A. Cancer in the Fourth Dimension: What Is the Impact of Circadian Disruption? Cancer Discov 10, 1455–1464 (2020). 10.1158/2159-8290.CD-20-0413

5 Kohsaka, A. et al. High-fat diet disrupts behavioral and molecular circadian rhythms in mice. Cell Metab 6, 414–421 (2007). 10.1016/j.cmet.2007.09.006

6 Panda, S. et al. Coordinated transcription of key pathways in the mouse by the circadian clock. Cell 109, 307–320 (2002). S0092867402007225 [pii]

7 Cox, K. H. & Takahashi, J. S. Circadian clock genes and the transcriptional architecture of the clock mechanism. J Mol Endocrinol 63, R93–R102 (2019). 10.1530/JME-19-0153

8 Ramsey, K. M. et al. Circadian clock feedback cycle through NAMPT-mediated NAD+ biosynthesis. Science 324, 651–654 (2009). 10.1126/science.1171641

9 Asher, G. et al. Poly(ADP-ribose) polymerase 1 participates in the phase entrainment of circadian clocks to feeding. Cell 142, 943–953 (2010). https://doi.org:S0092-8674(10)00940-2 [pii] 10.1016/j.cell.2010.08.016

10 Peek, C. B. et al. Circadian clock NAD+ cycle drives mitochondrial oxidative metabolism in mice. Science 342, 1243417 (2013). 10.1126/science.1243417

11 Yin, L. et al. Rev-erbalpha, a heme sensor that coordinates metabolic and circadian pathways. Science 318, 1786–1789 (2007). 1150179 [pii] 10.1126/science.1150179

12 Stokkan, K. A., Yamazaki, S., Tei, H., Sakaki, Y. & Menaker, M. Entrainment of the circadian clock in the liver by feeding. Science 291, 490–493 (2001).

13 Hepler, C. et al. Time-restricted feeding mitigates obesity through adipocyte thermogenesis. Science 378, 276–284 (2022). 10.1126/science.abl8007

14 Acosta-Rodriguez, V. A. et al. Misaligned feeding uncouples daily rhythms within brown adipose tissue and between peripheral clocks. Cell Rep 43, 114523 (2024). 10.1016/j.celrep.2024.114523

15 Hong, H. K. et al. Requirement for NF-kappaB in maintenance of molecular and behavioral circadian rhythms in mice. Genes Dev 32, 1367–1379 (2018). 10.1101/gad.319228.118

16 Paschos, G. K. et al. Obesity in mice with adipocyte-specific deletion of clock component Arntl. Nat Med 18, 1768–1777 (2012). 10.1038/nm.2979

17 Hepler, C. & Bass, J. Circadian mechanisms in adipose tissue bioenergetics and plasticity. Genes Dev 37, 454–473 (2023). 10.1101/gad.350759.123

18 Hasan, N. et al. Brown adipocyte-specific knockout of Bmal1 causes mild but significant thermogenesis impairment in mice. Mol Metab 49, 101202 (2021). 10.1016/j.molmet.2021.101202

19 Zhu, J., Vinothkumar, K. R. & Hirst, J. Structure of mammalian respiratory complex I. Nature 536, 354–358 (2016). 10.1038/nature19095

20 Stram, A. R. & Payne, R. M. Post-translational modifications in mitochondria: protein signaling in the powerhouse. Cell Mol Life Sci 73, 4063–4073 (2016). 10.1007/s00018-016-2280-4

21 Kleiner, S. et al. Development of insulin resistance in mice lacking PGC-1alpha in adipose tissues. Proc Natl Acad Sci U S A 109, 9635–9640 (2012). 10.1073/pnas.1207287109

22 Chouchani, E. T. & Kajimura, S. Metabolic adaptation and maladaptation in adipose tissue. Nat Metab 1, 189–200 (2019). 10.1038/s42255-018-0021-8

23 Bogacka, I., Xie, H., Bray, G. A. & Smith, S. R. Pioglitazone induces mitochondrial biogenesis in human subcutaneous adipose tissue in vivo. Diabetes 54, 1392–1399 (2005). 10.2337/diabetes.54.5.1392

24 Semple, R. K. et al. Expression of the thermogenic nuclear hormone receptor coactivator PGC-1alpha is reduced in the adipose tissue of morbidly obese subjects. Int J Obes Relat Metab Disord 28, 176–179 (2004). 10.1038/sj.ijo.0802482

25 McElroy, G. S. et al. NAD+ Regeneration Rescues Lifespan, but Not Ataxia, in a Mouse Model of Brain Mitochondrial Complex I Dysfunction. Cell Metab 32, 301–308 e306 (2020). 10.1016/j.cmet.2020.06.003

26 Han, S. et al. Mitochondrial integrated stress response controls lung epithelial cell fate. Nature 620, 890–897 (2023). 10.1038/s41586-023-06423-8

27 Seo, B. B. et al. Molecular remedy of complex I defects: rotenone-insensitive internal NADH-quinone oxidoreductase of Saccharomyces cerevisiae mitochondria restores the NADH oxidase activity of complex I-deficient mammalian cells. Proc Natl Acad Sci U S A 95, 9167–9171 (1998). 10.1073/pnas.95.16.9167

28 Park, J. S., Li, Y. F. & Bai, Y. Yeast NDI1 improves oxidative phosphorylation capacity and increases protection against oxidative stress and cell death in cells carrying a Leber’s hereditary optic neuropathy mutation. Biochim Biophys Acta 1772, 533–542 (2007). 10.1016/j.bbadis.2007.01.009

29 Wen, H. et al. Mitochondrial diseases: from molecular mechanisms to therapeutic advances. Signal Transduct Target Ther 10, 9 (2025). 10.1038/s41392-024-02044-3

30 Fernandez-Aguera, M. C. et al. Oxygen Sensing by Arterial Chemoreceptors Depends on Mitochondrial Complex I Signaling. Cell Metab 22, 825–837 (2015). 10.1016/j.cmet.2015.09.004

31 Tuppen, H. A. et al. The p.M292T NDUFS2 mutation causes complex I-deficient Leigh syndrome in multiple families. Brain 133, 2952–2963 (2010). 10.1093/brain/awq232

32 Enguix, N. et al. Mice lacking PGC-1beta in adipose tissues reveal a dissociation between mitochondrial dysfunction and insulin resistance. Mol Metab 2, 215–226 (2013). 10.1016/j.molmet.2013.05.004

33 Geoghegan, G. et al. Targeted deletion of Tcf7l2 in adipocytes promotes adipocyte hypertrophy and impaired glucose metabolism. Mol Metab 24, 44–63 (2019). 10.1016/j.molmet.2019.03.003

34 Garritson, J. D. et al. BMPER is a marker of adipose progenitors and adipocytes and a positive modulator of adipogenesis. Commun Biol 6, 638 (2023). 10.1038/s42003-023-05011-w

35 Lazarou, M., Thorburn, D. R., Ryan, M. T. & McKenzie, M. Assembly of mitochondrial complex I and defects in disease. Biochim Biophys Acta 1793, 78–88 (2009). 10.1016/j.bbamcr.2008.04.015

36 Nakahata, Y., Sahar, S., Astarita, G., Kaluzova, M. & Sassone-Corsi, P. Circadian control of the NAD+ salvage pathway by CLOCK-SIRT1. Science 324, 654–657 (2009). 10.1126/science.1170803

37 Cela, O. et al. Clock genes-dependent acetylation of complex I sets rhythmic activity of mitochondrial OxPhos. Biochim Biophys Acta 1863, 596–606 (2016). 10.1016/j.bbamcr.2015.12.018

38 Peek, C. B. et al. Circadian Clock Interaction with HIF1alpha Mediates Oxygenic Metabolism and Anaerobic Glycolysis in Skeletal Muscle. Cell Metab 25, 86–92 (2017). 10.1016/j.cmet.2016.09.010

39 Timmons, G. A. et al. The Circadian Clock Protein BMAL1 Acts as a Metabolic Sensor In Macrophages to Control the Production of Pro IL-1beta. Front Immunol 12, 700431 (2021). 10.3389/fimmu.2021.700431

40 Alexander, R. K. et al. Bmal1 integrates mitochondrial metabolism and macrophage activation. Elife 9 (2020). 10.7554/eLife.54090

41 Zhu, P. et al. BMAL1 drives muscle repair through control of hypoxic NAD(+) regeneration in satellite cells. Genes Dev 36, 149–166 (2022). 10.1101/gad.349066.121

42 Krycer, J. R. et al. Lactate production is a prioritized feature of adipocyte metabolism. J Biol Chem 295, 83–98 (2020). 10.1074/jbc.RA119.011178

43 TeSlaa, T. et al. The Source of Glycolytic Intermediates in Mammalian Tissues. Cell Metab 33, 367–378 e365 (2021). 10.1016/j.cmet.2020.12.020

44 Chandel, N. S. Mitochondria as signaling organelles. BMC Biol 12, 34 (2014). 10.1186/1741-7007-12-34

45 Sarvari, A. K. et al. Plasticity of Epididymal Adipose Tissue in Response to Diet-Induced Obesity at Single-Nucleus Resolution. Cell Metab 33, 437–453 e435 (2021). 10.1016/j.cmet.2020.12.004

46 Emont, M. P. et al. A single-cell atlas of human and mouse white adipose tissue. Nature 603, 926–933 (2022). 10.1038/s41586-022-04518-2

47 Salabei, J. K., Gibb, A. A. & Hill, B. G. Comprehensive measurement of respiratory activity in permeabilized cells using extracellular flux analysis. Nat Protoc 9, 421–438 (2014). 10.1038/nprot.2014.018

